# Janusein and EB Cooperate to Enhance Microtubule Plus-end Catastrophe

**DOI:** 10.64898/2026.10.26.761828

**Authors:** Kaiming Xu, Zhengyang Guo, Yishuo Cao, Yuqi Ye, Cuirong Guan, Kai Jiang, Wei Li, Guangshuo Ou

**Affiliations:** State Key Laboratory of Membrane Biology, Tsinghua-Peking Center for Life Sciences, Beijing Frontier Research Center for Biological Structure, McGovern Institute for Brain Research, MOE Key Laboratory for Protein Science, School of Life Sciences, Tsinghua University, Beijing, 100084, China; School of Basic Medical Sciences, Institute of Biomedical Innovation, Jiangxi Medical College, Nanchang University, Nanchang, 330031, China; State Key Laboratory of Technologies for Chinese Medicine Pharmaceutical Process Control and Intelligent Manufacture, Nanjing University of Chinese Medicine, Nanjing, 210023, China; Frontier Science Center for Immunology and Metabolism, Medical Research Institute, Wuhan University, Wuhan, 430072, China; State Key Laboratory of Membrane Biology, School of Basic Medical Sciences, Tsinghua University, Beijing, 100084, China

## Abstract

Microtubule (MT) plus ends are organized by end-binding (EB) proteins, yet how EB-associated factors modulate the transition from growth to catastrophe remains incompletely understood. Using spatially resolved proximity proteomics, we identified Janusein (ZC2HC1A) as a conserved, poorly characterized MT-associated protein (MAP). At low expression levels, Janusein tracked MT plus ends and reduced EB comet velocity; at higher levels, it redistributed to the MT lattice and was associated with MT bundling and widespread MT loss. Loss of Janusein resulted in spindle defects in human cells. In vitro, Janusein alone facilitated MT elongation through its C-terminal fragment, but promoted MT catastrophe through its N-terminal fragment in the presence of EB proteins. Molecular dynamics simulations and mutational analyses suggest that an N-terminal loop may perturb lateral contacts between adjacent MT protofilaments. Together, these findings identify Janusein as an unconventional MAP that functionally cooperates with EB proteins to promote MT catastrophe.

## Introduction

Microtubules (MTs) are dynamic polymers that stochastically switch between phases of growth and shrinkage, a behavior termed ‘dynamic instability’ (Brouhard and Rice, 2018; McIntosh, 2025; Mitchison and Kirschner, 1984). Dynamic instability enables MT arrays to be rapidly remodeled in response to changing cellular demands such as spindle assembly and axon morphogenesis (Kraus et al., 2023; Travis et al., 2022; Valdez et al., 2023; Yogev et al., 2016).

MT dynamics are shaped by tubulin isotypes and post-translational modifications, as well as by a diverse repertoire of microtubule-associated proteins (MAPs) that regulate MT polymerization, stability, organization, and turnover (Bodakuntla et al., 2019; Goodson and Jonasson, 2018; Janke and Magiera, 2020; McKenna et al., 2023; Roll-Mecak, 2020). While many MAPs stabilize MTs—such as Tau in neurons or CLASPs at the cortex (Akhmanova and Kapitein, 2022; Bravo et al., 2024)—the controlled disassembly of MTs is equally crucial for turnover and network remodeling. Several mechanistically distinct pathways promote MT disassembly. The AAA ATPases katanin and spastin sever MTs by using ATP hydrolysis to extract tubulin subunits from the lattice, generating new MT ends that can subsequently depolymerize or regrow (Hartman et al., 1998; McNally and Vale, 1993; Roll-Mecak and Vale, 2008; Vemu et al., 2018; Zehr et al., 2017). Kinesin-13 and kinesin-8 motors induce protofilament peeling at MT ends through nucleotide-driven conformational cycling (Benoit et al., 2018; Desai et al., 1999; Gardner et al., 2011; Gupta et al., 2006; Hunter et al., 2003; Varga et al., 2009). Moreover, stathmin and the tubulin-binding PN2-3 domain of CPAP can inhibit MT assembly by sequestering soluble tubulin heterodimers, thereby reducing the pool of assembly-competent tubulin in living organisms (Belmont and Mitchison, 1996; Cormier et al., 2009; Ravelli et al., 2004; Tang et al., 2009).

A specialized subset of MAPs, termed plus-end-tracking proteins (+TIPs), preferentially accumulates at growing MT plus ends and regulates MT dynamics (Akhmanova and Kapitein, 2022; Akhmanova and Steinmetz, 2015; Gudimchuk and McIntosh, 2021; Janke and Magiera, 2020; McIntosh, 2025; Paz and Lüders, 2018; Wu and Akhmanova, 2017). +TIPs form a dynamic interaction network at growing MT ends, with end-binding (EB) proteins serving as core components of this network (Akhmanova and Steinmetz, 2015; Bieling et al., 2007; Miesch et al., 2023; Saunders et al., 2025). EBs autonomously recognize a nucleotide-dependent structural state of the growing MT lattice and form characteristic comet-like distributions behind the plus end of the growing MT (Maurer et al., 2014; Zhang et al., 2015). Through their C-terminal interaction region, EBs recruit diverse +TIPs—including CLIP-170, CLASPs, and other SxIP motif-containing proteins—that regulate MT dynamics in cooperation (Akhmanova and Steinmetz, 2015; Brouhard et al., 2008; Reber et al., 2013; Slep and Vale, 2007; Vitre et al., 2008). Moreover, studies reveal that EB and associated +TIPs can form multivalent, condensate-like assemblies at plus ends that may potentiate complex regulatory activities (Maan et al., 2023; Meier et al., 2023; Song et al., 2023).

The effects of EB proteins on MT plus-end dynamics are context dependent. Mammalian EB proteins, as well as the fission-yeast EB ortholog Mal3, increase MT catastrophe frequency in vitro (Bieling et al., 2007; Vitre et al., 2008; Zanic et al., 2013). By contrast, EB proteins promote persistent MT growth by suppressing catastrophes in cultured mammalian cells (Komarova et al., 2009). Therefore, the molecular basis of these context-dependent effects remains incompletely understood, and the molecular determinants that switch EB networks between growth-promoting and catastrophe-promoting states remain poorly defined.

In this study, we identified ZC2HC1A (Janusein), an evolutionarily conserved MAP that promotes MT catastrophe in synergy with EB proteins. Its N-terminal fragment induces catastrophe, while the C-terminal fragment stabilizes and bundles MTs. Named after the two-faced Roman god Janus, Janusein exhibits context-dependent activities in regulating MT dynamics. Computational and mutational analyses suggest that the N-terminal fragment may perturb lateral contacts between adjacent MT protofilaments. Janusein depletion in HeLa cells caused mitotic spindle defects, and a *C. elegans* suppressor screen revealed that loss of the Janusein homologue rescues ciliary defects caused by a mutant α-tubulin, supporting a physiological role for Janusein homologues in MT regulation. Together, these findings reveal context-dependent activities of Janusein in MT regulation and functional cooperation between Janusein and EB proteins in promoting catastrophe.

## Results

### Proximity proteomics and AlphaFold3-assisted interface screening identify Janusein as a microtubule-associated protein

To identify MAPs with the potential to bind the MT lattice, we combined spatially resolved proximity proteomics with structure-based interface screening. We developed a minimally perturbative split-TurboID labeling strategy, adapted from split-GFP tubulin tagging, that enables selective biotinylation of proteins at either the luminal or cytoplasmic surface of MTs in living cells (Figures 1A-G; Figures S1A-E) (Cho et al., 2020a; Cho et al., 2020b; Xu et al., 2024). The TurboID N-terminal fragment was inserted into the lumen-facing H1-S2 loop of α-tubulin (TUBA4A) or fused to the C terminus of β-tubulin (TUBB8) (Figures 1C-D), two sites previously shown to tolerate tagging with minimal impact on MT behavior (Feng et al., 2016; Nogales et al., 1998; Schatz et al., 1987). Quantitative proteomic analysis in HeLa and HEK293T cells identified 132 proteins reproducibly enriched across both datasets (Figure S2; Table S1-S2; see Methods). Western blot analyses validated selective enrichment of the known luminal MAP (JPT1) (Shao et al., 2026), and an outer-surface MAP (EML4) in their respective fractions (Figure S2D) (Jijumon et al., 2022).

**Figure 1.**
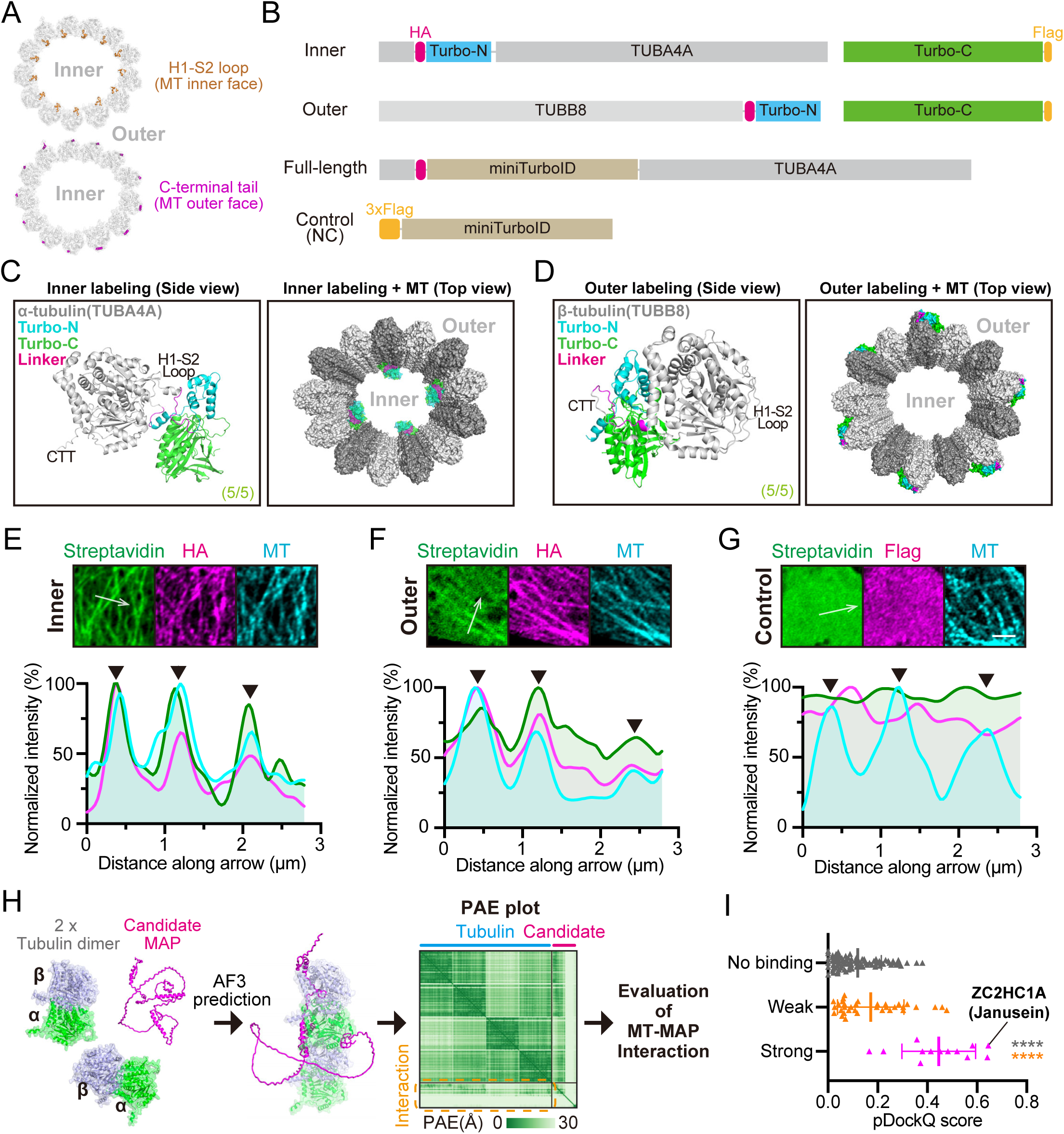
Identification of MT-associated proteome by proximity labeling and AlphaFold3-enabled interaction prediction. (A) Structural models showing localizations of H1-S2 loop (brown) and C-terminal tail (magenta) on the luminal and outer surfaces of MTs, respectively. The 13-protofilament MT model (PDB ID: 6U42) was employed as the template. (B) Linear representation of protein constructs for inner proximity labeling, outer proximity labeling, full-length labeling and negative control (NC). For inner, an HA tag and split-Turbo-N were inserted into the H1-S2 loop of TUBA4A. Turbo-C was provided separately. For outer, an HA tag and split-Turbo-N were fused to the C-terminus of TUBB8. (C and D) AlphaFold3-predicted multimer structures of designed inner (C) and outer (D) constructs. For each prediction, all 5 independent cases showed the same results (5/5). Predicted structures were then aligned to tubulins in a MT structural model (PDB: 6U42). The recombinant split-TurboID enzyme (cyan and green) located within the MT lumen (inner), or on the MT outer surface (outer). CTT, C-terminal tail. (E-G) (Top) Representative immunofluorescence images showing differential biotinylation patterns in inner (E), outer (F), and control (G) labeling systems in HeLa cells. Streptavidin (green) marks the biotinylated proteome in cells. HA or Flag tag labels the engineered TurboID constructs. Scale bar, 2 μm. (Bottom) Measured fluorescence intensities along the white arrows in each group. Maximal intensities are normalized to 100%. Black arrowheads indicate the positions of MTs. (H) Schematic model illustrating AlphaFold3 (AF3)-enabled prediction of MT-MAP interaction patterns. Enriched candidate MAPs identified in both HeLa and HEK293T cells (N = 132) were subjected to prediction. The affinity of each MAP to the MT protofilament is assessed according to the interaction regions (as reflected by the green zone, within the orange dashed box) within the PAE (predicted alignment error) plot. Expected position error (Å) ranges from 0 (green) to 30 (white). In this schematic, the representative MAP (magenta) is ZC2HC1A (Janusein). (I) Predicted DockQ (pDockQ) scores for all AlphaFold3-predicted MT-MAP interactions (N = 132; from panel H). ZC2HC1A (Janusein) displays the highest pDockQ score. Shown as mean ± SD. Nonparametric tests (Kruskal-Wallis test with Dunn’s multiple comparisons) were performed. **** p < 0.0001.

Because proximity labeling does not distinguish direct lattice binding from indirect association, we next evaluated whether these candidates could engage MTs through direct physical interfaces. Each protein was modeled using AlphaFold3 together with two α/β-tubulin heterodimers arranged as a short protofilament, providing a standardized structural context for assessing MT interactions (Figure 1H). Predicted alignment error (PAE) at the interface and predicted DockQ (pDockQ) scores were used to estimate interaction confidence (Bryant et al., 2022). On the basis of PAE metrics, candidates segregated into strong, weak, or no predicted MT-binding classes (Figure S2G), with strong-binding models exhibiting significantly higher pDockQ scores and exceeding established confidence thresholds (Figure 1I). Among all candidates, a previously uncharacterized zinc-finger protein ZC2HC1A—hereafter termed Janusein—stood out by displaying the highest predicted interface confidence (pDockQ=0.65) (Figure 1I; Table S3). This strong and recurrent predicted interaction prioritized Janusein for subsequent experimental and mechanistic characterization.

### Janusein exhibits dose-dependent plus-end tracking and microtubule destabilization in cells

To define how Janusein influences MT organization, we first examined its behavior across a range of expression levels. Janusein displayed concentration-dependent activities (Figure S3A). At low expression, EGFP-Janusein formed comet-like puncta that co-migrated with EB3-mCherry at growing MT plus ends, demonstrating their co-localization (Figures 2A-B; Movie S1). Even in this low-expression regime, Janusein perturbed MT dynamics, reducing polymerization rates and shifting the growth-shrinkage balance toward instability (Figures 2B-C). As expression levels increased, Janusein redistributed from plus ends onto the MT lattice, where it partially decorated MTs and promoted bundling (Figures 2D-F). At high expression, Janusein induced widespread MT loss and accumulated in cytoplasmic condensates (Figure 2G), revealing potent dose-dependent destabilizing activity (Figure S3B).

**Figure 2.**
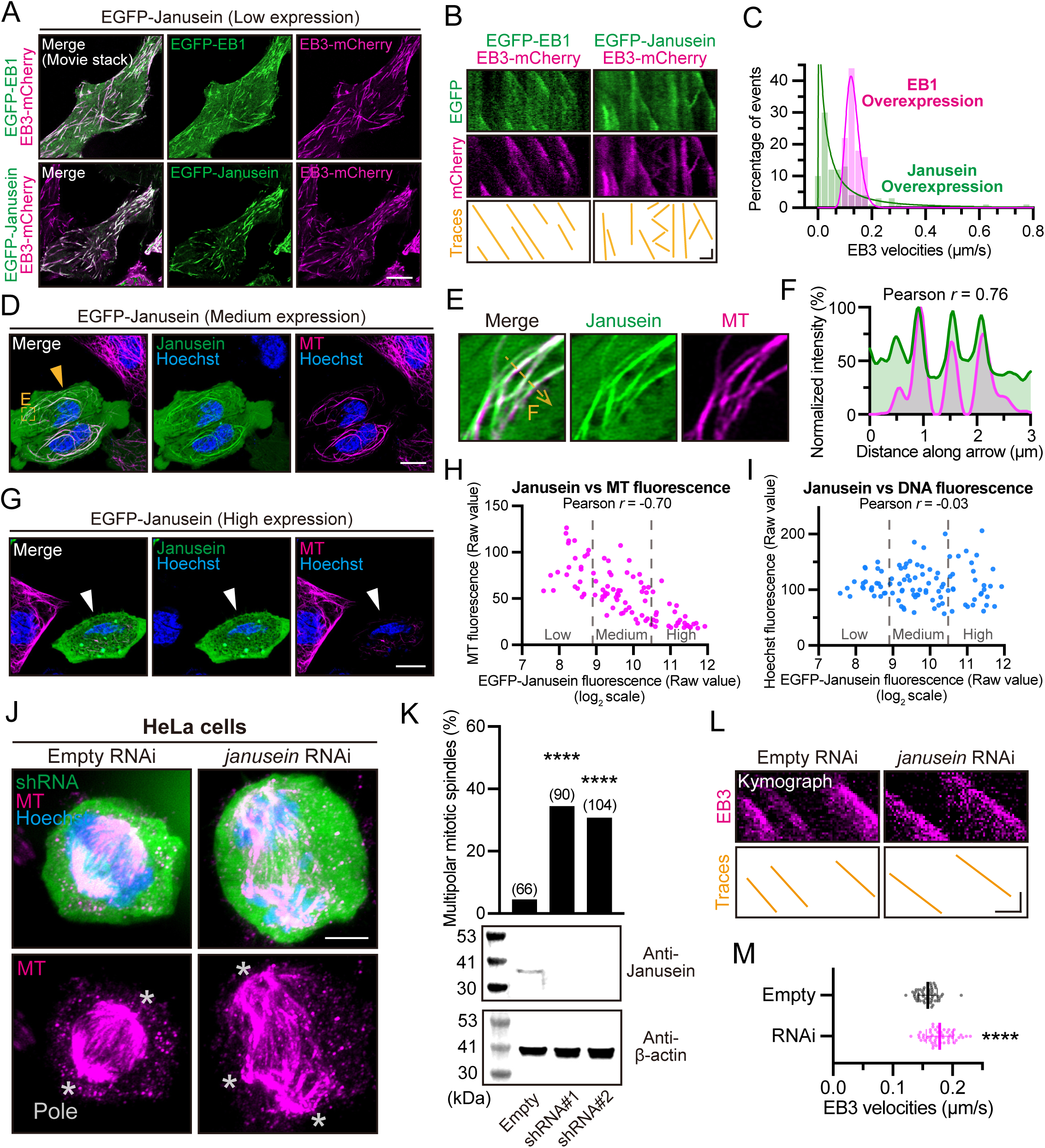
Janusein is a MT plus-end-binding protein with dose-dependent activities. (A) Representative live images of HeLa cells with low expression of EGFP-EB1 and EB3-mCherry (Left), or EGFP-Janusein and EB3-mCherry (Right). Image stacks spanning 20 sec are superimposed for visualization. See also Movie S1. Scale bar, 10 μm. (B) Representative kymographs showing movements of EB1, EB3 or Janusein in HeLa cells. Scale bar, 2 μm (horizontal) and 10 sec (vertical). (C) Distribution of EB3 velocities in EB1 & EB3 overexpression, or Janusein & EB3 overexpression cells. N = 50 (EB1) or 100 (Janusein) moving events. (D) Representative live-cell images of HeLa cells with medium expression of EGFP-Janusein. Transfection-positive cells are indicated by orange arrowhead. Tubulin Tracker Deep Red (magenta) was used to stain MT networks. Scale bar, 10 μm. (E) Enlarged view showing co-localization of Janusein (green) and MTs (magenta) in (D). (F) Fluorescence distribution along the arrow in (E). The maximal fluorescence intensities are normalized to 100%. (G) Representative live-cell images of HeLa cells with high expression of EGFP-Janusein. Transfection-positive cells are indicated by white arrowhead. Scale bar, 10 μm. (H and I) Correlation scatter plots of whole-cell EGFP-Janusein fluorescence (horizontal axis) and MT fluorescence (H, vertical axis) or DNA fluorescence (I, vertical axis). Raw fluorescence values are measured and presented. Cells with low expression (Raw value < 500), medium expression (500 ≤ Raw value ≤ 1500), or high expression (Raw value > 1500) of EGFP-Janusein are distinguished by gray dashed lines. N = 100 cells. (J) Representative immunofluorescence images of shRNA-CopGFP-transfected mitotic HeLa cells. Spindle poles are indicated by asterisks. Scale bar, 5 μm. (K) (Top) Percentage of shRNA-positive cells with multipolar spindles. Metaphase HeLa cells were subjected for quantification. Chi-Square tests are performed. (Bottom) Western blot analysis showing Janusein or β-actin levels in HeLa total lysates after treatments with empty shRNA, Janusein shRNA#1 or shRNA#2. (L) Representative kymographs showing EB3 traces in Janusein shRNA-positive cells. Scale bar, 2 μm (horizontal) and 10 sec (vertical). (M) Quantification of EB3 velocities in empty, or Janusein RNAi cells. Shown as mean ± SD. Unpaired t tests (Welch’s correction) were performed. N = 50 moving events. **** p < 0.0001.

Quantitative analysis demonstrated a strong negative correlation between EGFP-Janusein intensity and MT fluorescence (Pearson’s *r* = -0.70) (Figures 2H-I), supporting the relationship between Janusein abundance and MT loss. In contrast, overexpression of EB1 or EB3 did not trigger MT depolymerization under comparable conditions (Figures S3C-D). Although EB proteins can undergo phase separation in vitro (Maan et al., 2023; Meier et al., 2023; Song et al., 2023), they rarely form prominent condensates in cells (Figures S3C and S3E), underscoring the distinct cellular behavior of Janusein. These data identify Janusein as an unconventional MAP whose activity shifts from plus-end tracking to lattice association and ultimately to robust MT destabilization as its concentration rises.

### Janusein is required for spindle integrity and restrains microtubule growth

To determine Janusein’s physiological role, we depleted endogenous Janusein in HeLa cells using two independent shRNAs (Figures 2J-K). Loss of Janusein resulted in a marked increase in multipolar spindles during mitosis compared with control cells (Figures 2J-K), indicating defective spindle organization. These phenotypes resemble defects observed upon disruption of established MT-regulating factors required for spindle assembly (Ganem and Compton, 2004; McNally et al., 2006), indicating that Janusein contributes to mitotic MT remodeling. Consistent with the reduced MT growth rate observed upon Janusein overexpression (Figure 2C), Janusein depletion significantly increased MT growth rate in interphase cells, as measured by EB comet tracking (Figures 2L-M). Live-cell imaging of a stable EGFP-Janusein cell line revealed cell cycle-dependent redistribution (Figure S4A). Janusein tracked MT plus ends during interphase and relocalized broadly across the mitotic spindle during metaphase (Figure S4A), suggesting direct engagement with spindle MTs.

### Genetic suppression of ciliary defects in *C. elegans* by Janusein homolog

Janusein expression was detected across diverse human tissues (Figure S4B), and phylogenetic analysis identified conserved homologs throughout eukaryotes, from algae to mammals (Figures S4C-D), supporting a conserved role in regulating eukaryotic MTs. We also performed a genetic suppressor screen in *C. elegans* (see Methods) to identify suppressors of ciliary defects caused by the α-tubulin TBA-5 (A19V) mutation which disrupts ciliary MT architecture (Figure S4E) (Hao et al., 2011; Xu et al., 2026b; Xu et al., 2024). A putative null allele of the Janusein homolog, ZCHC-1A, was identified as a strong suppressor, rescuing the ciliary defects in TBA-5 (A19V) mutants and restoring ciliary MT length (Figures S4F-G). Importantly, re-expression of GFP-tagged wild-type ZCHC-1A in *tba-5 (A19V); zchc-1A (null)* double mutants abolished this suppression, resulting in defective cilia mirroring those in *tba-5 (A19V)* single mutants (Figures S4F-G). While the deletion of ZCHC-1A rescued ciliary defects (Figure S4H), it also reduced the brood size of *C. elegans* to approximately 80% of wild-type levels (Figure S4I), suggesting potential defects in the germline or early embryos. These findings further highlight the in vivo function of Janusein homologs in MT regulation in organismal levels and across species.

### Distinct Janusein domains mediate microtubule destabilization and stabilization

AlphaFold-based modeling predicted that Janusein comprises an N-terminal folded domain (NTD), a central intrinsically disordered region (IDR), and a structured C-terminal domain (CTD) (Figures 3A-B). To define how these regions contribute to Janusein activity, we expressed EGFP-tagged truncation mutants in HeLa cells (Figures S5A-B). At expression levels that cause robust MT loss by full-length Janusein, the isolated NTD was sufficient to disassemble MT networks, whereas deletion of the NTD abolished MT loss despite high expression (Figures 3C-D). Removal of either the IDR or CTD did not impair NTD-driven MT loss, identifying the NTD as the core destabilizing module (Figures 3C-D). Immunofluorescence corroborated these observations (Figure S5C). Under these conditions, Janusein condensates did not visibly sequester cytoplasmic tubulin pools (Figures S5D-E). In contrast, expression of the IDR-CTD fragment induced extensive MT bundling and lattice stabilization (Figure 3C), generating MT arrays resistant to drug-induced MT loss (Figures S5F-G).

**Figure 3.**
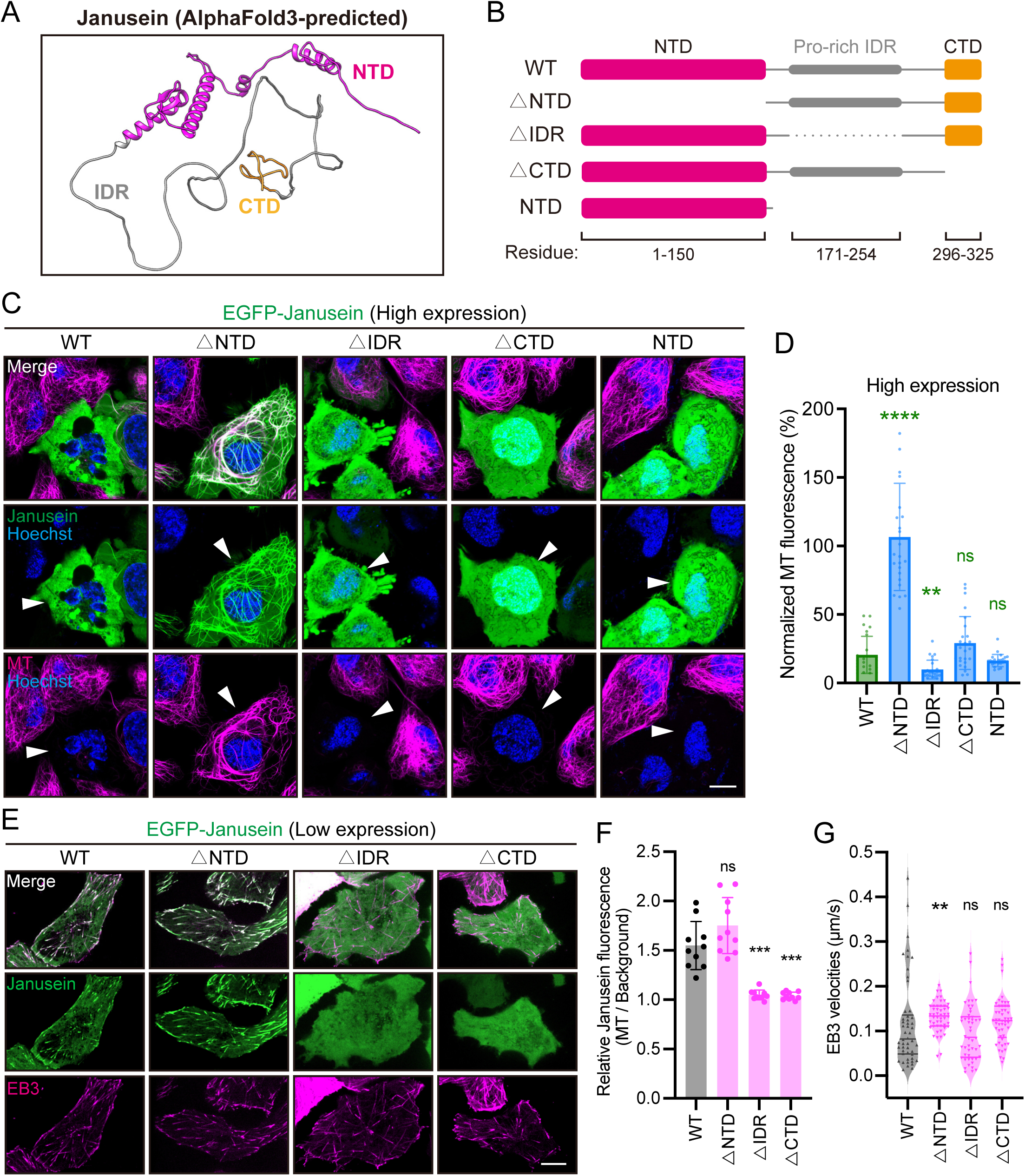
Functional dissection of Janusein domains. (A) AlphaFold3-predicted structure of Janusein. The N-terminal domain (NTD) of Janusein is colored in magenta, intrinsically disordered region (IDR) in gray, and the C-terminal domain (CTD) in orange. (B) Schematic of Janusein truncation mutants. (C) Representative live images of HeLa cells with high expression of EGFP-Janusein truncation mutants. Transfection-positive cells are indicated by white arrowheads. Tubulin Tracker Deep Red (magenta) was used to stain MT networks. Scale bar, 10 μm. (D) Whole-cell MT fluorescence intensities in HeLa cells with high expression of Janusein truncation mutants. For each group, the average MT fluorescence of 10 randomly selected transfection-negative cells is normalized to 100%. Shown as mean ± SD. Welch’s ANOVA with Dunnett’s tests were performed. N = 25 cells. ns, not significant; *** p < 0.001; **** p < 0.0001. (E) Representative live images of HeLa cells with low expression of EB3-mCherry and EGFP-Janusein truncation mutants. Image stacks spanning 20 sec are super-imposed for visualization. Scale bar, 10 μm. (F) EGFP-Janusein fluorescence ratio of MTs to cytosol backgrounds in each group. N = 10 cells. For each cell, the average ratio of 10 MTs to their respective background fluorescence is calculated. Shown as mean ± SD. Welch’s ANOVA with Dunnett’s tests were performed. (G) EB3 velocities in each group. N = 50 moving events. Shown as median ± IQR (interquartile range). Kruskal-Wallis test with Dunn’s multiple comparisons were performed.

At low expression levels, where full-length Janusein behaves as a plus-end-tracking protein, the domain requirements differed. Deletion of the NTD minimally affected plus-end association, whereas removal of either the IDR or CTD markedly reduced MT binding (Figures 3E-G). Mutations of conserved residues within the CTD weakened lattice association, confirming a direct MT-binding interface (Figures S5H-J). Consistent with these observations, deletion of the NTD restored MT growth rates reduced by full-length Janusein expression (Figure 3G). These data support a model in which Janusein is a modular protein with an N-terminal destabilizing module counterbalanced by a C-terminal MT-binding and stabilizing module, enabling expression-dependent switching between growth restraint, lattice stabilization, and MT disassembly.

### Micromolar Janusein promotes microtubule bundling and suppresses catastrophe in vitro

To determine whether Janusein directly destabilizes MTs, we reconstituted MT dynamics in vitro using purified Janusein (Figures S6A-C). Contrary to cellular MT-loss phenotype associated with Janusein overexpression, full-length Janusein alone did not affect in vitro MT growth at nanomolar concentrations (250 nM) (Figures 4A-B). However, at micromolar concentrations (4 μM), it markedly reduced catastrophe frequency and increased maximal growth length without significantly altering MT growth rate (Figures 4A-B). Truncation analysis revealed that the IDR-CTD, rather than the NTD, mediated apparent MT association and reduced catastrophe frequency (Figures 4A-C). When tested at 4 μM, full-length Janusein and the IDR-CTD, but not the NTD, promoted lateral association of MTs, producing bundled filaments during elongation (Figures S6D-E). Together, these findings indicate that the Janusein IDR-CTD mediates MT association, bundling, and stabilization under the 4 μM conditions tested in vitro, consistent with the MT bundling and stabilization observed in cells (Figures 3C and S5F-G).

**Figure 4.**
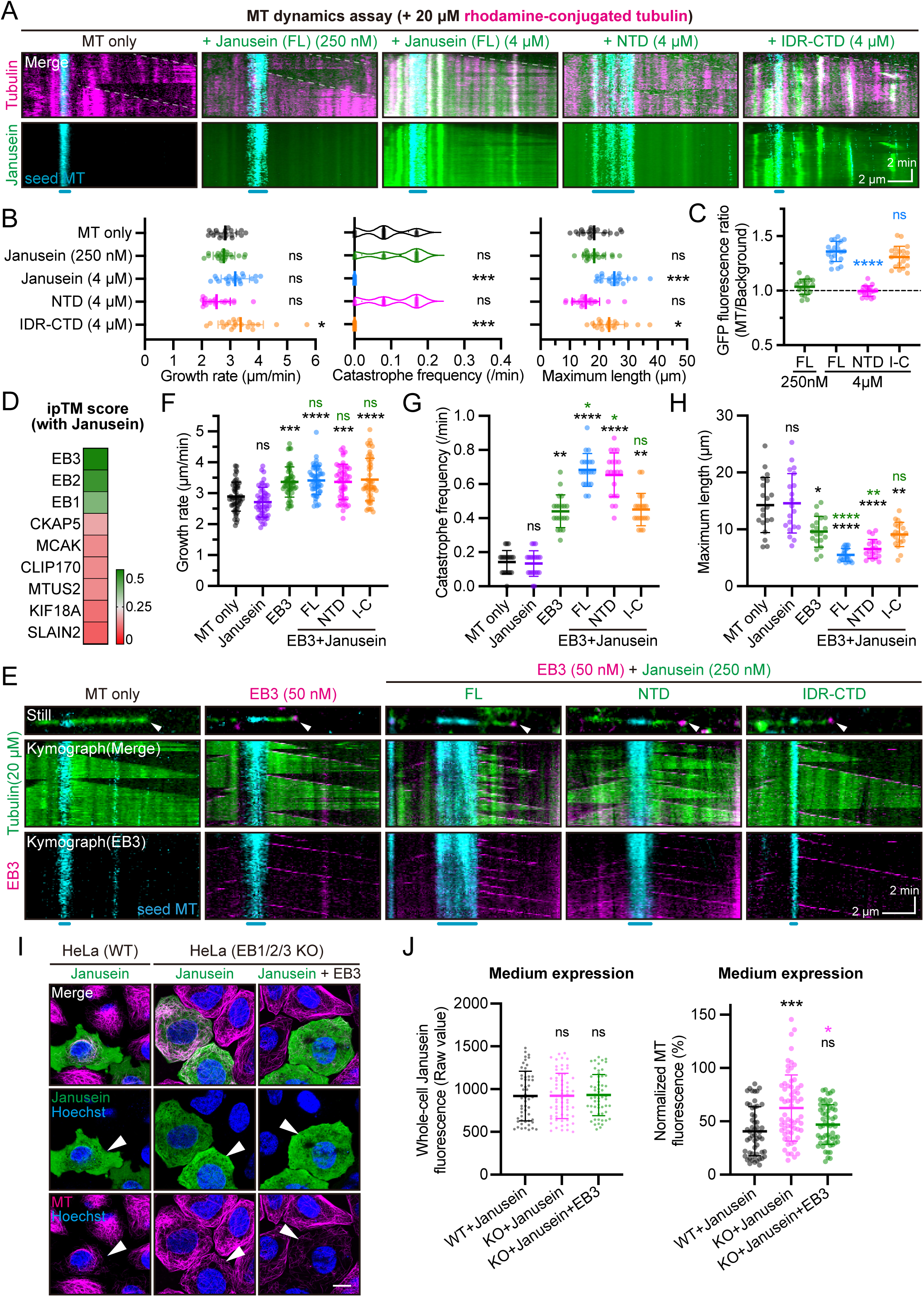
Janusein cooperates with EB family proteins to destabilize MTs in vitro. (A) Representative kymographs displaying the time-lapse dynamics of growing MTs. HiLyte647-conjugated seed MTs (cyan), rhodamine-conjugated tubulins (magenta), and EGFP-Janusein (green) were employed (see Methods). White dashed lines within kymographs mark the plus ends of seed MTs where dynamic MTs grow out. Scale bar, 2 μm (horizontal) and 2 min (vertical). (B) MT growth rate (left), catastrophe frequency (middle), and maximal growth length (right) for each group in (A) (N = 20). One-way ANOVA with Dunnett’s tests (left, right), or Kruskal-Wallis test with Dunn’s multiple comparisons (middle) were performed. (C) Fluorescence ratios of GFP-Janusein on dynamic MTs relative to adjacent cytosolic background (N = 20). Shown as mean ± SD. One-way ANOVA with Dunnett’s tests were performed. (D) Heatmap representation of ipTM (interface predicted template modeling) scores of Janusein with nine representative +TIPs as predicted by AlphaFold3. ipTM scores range from 0 (red) to 0.5 (green). See also Figure S6G for detailed PAE (predicted alignment error) plots. (E) Representative stills and kymographs displaying the time-lapse dynamics of growing MTs. HiLyte647-conjugated seed MTs (cyan), HiLyte488-conjugated tubulins (green), EGFP-Janusein (green), and EB3-mCherry (magenta) were employed. White arrowheads indicated dynamic MT plus ends. Scale bar, 2 μm (horizontal) and 2 min (vertical). See also Movie S2. (F-H) MT growth rate (F) (N = 50), catastrophe frequency (G) (N = 20), and maximal growth length (H) (N = 20) for each group in (E). One-way ANOVA with Dunnett’s tests (F, H), or Kruskal-Wallis test with Dunn’s multiple comparisons (G) were performed. (I) Representative live images of wild-type (WT) HeLa cells or EB1/2/3 triple-knockout (KO) HeLa cells with medium expression of EGFP-Janusein alone, or EGFP-Janusein + EB3-mCherry. Transfection-positive cells are indicated by white arrowheads. Tubulin Tracker Deep Red (magenta) was used to stain MT networks. Scale bar, 10 μm. (J) Whole-cell EGFP-Janusein (left) or MT (right) fluorescence intensities in the same batch of HeLa cells (N > 50 cells) with medium expression of Janusein (Raw value of GFP fluorescence: 500-1500). For MT fluorescence intensities (right), the average whole-cell MT fluorescence of 20 randomly selected transfection-negative cells is normalized to 100% in each group. Shown as mean ± SD. One-way ANOVA with Dunnett’s tests were performed. ns, not significant; * p < 0.05; ** p < 0.01; *** p < 0.001; **** p < 0.0001.

Furthermore, Janusein lacks recognizable ATPase motifs. At 6 μM, it bound to GMPCPP-stabilized MTs but failed to sever, fragment, or depolymerize the lattice in the presence of ATP (Figure S6F), distinguishing it from classical ATP-dependent severing enzymes (Hartman et al., 1998; Roll-Mecak and Vale, 2008). Thus, Janusein alone stabilizes and bundles MTs in vitro; these activities do not explain Janusein-associated MT loss in cells, suggesting that its MT-destabilizing activity requires additional cellular components.

### Janusein cooperates with EB to induce MT catastrophe through its NTD

Because Janusein tracks growing MT plus ends, we hypothesized that plus-end-associated proteins may enable its destabilizing activity. To identify candidate plus-end-associated partners, we first performed AlphaFold3-based interaction screening of Janusein with a panel of canonical +TIPs (Bieling et al., 2007; Brouhard et al., 2008; Honnappa et al., 2009; Jiang et al., 2009; Perez et al., 1999; Song et al., 2023; van der Vaart et al., 2011). EB family proteins emerged as the strongest predicted partners, with EB3 exhibiting the highest interface confidence (Figures 4D and S6G). To test this prediction, we reconstituted MT dynamics in vitro with purified Janusein (250 nM) and EB3 (50 nM) (Figure 4E). EB3 alone increased both MT growth rate and catastrophe frequency, as expected (Figures 4F-G; Movie S2) (Bieling et al., 2007). However, combining EB3 with full-length (FL) Janusein further promoted MT catastrophe without affecting growth rates, producing a marked increase in depolymerization events relative to either protein alone (Janusein: 0.13 ± 0.08 min⁻¹; EB3: 0.44 ± 0.10 min⁻¹; FL + EB3: 0.68 ± 0.10 min⁻¹) (Figures 4E-H; Movie S2). Domain reconstitution demonstrated that this synergistic effect was mediated by the Janusein N-terminal domain: the NTD plus EB3 was sufficient to induce catastrophe, whereas the IDR-CTD exhibited only minimal activity beyond that of EB3 alone (Figures 4E-H; Movie S2). This effect is dose-dependent with respect to Janusein concentration (Figures S6H-I), further supporting a direct role of Janusein in MT depolymerization. Notably, full-length Janusein effectively promoted MT catastrophe at a physiologically relevant concentration (50 nM) in the presence of EB3 (Figures S6H-I), suggesting that the observed in vitro activity is mechanistically coupled to its cellular function.

We next asked whether this functional synergy between EB proteins and Janusein is recapitulated in cells (Figure 4I). Whereas full-length Janusein induced pronounced MT loss in wild-type HeLa cells, this activity was markedly attenuated in EB1/2/3 triple-knockout cells despite comparable Janusein expression levels (Figures 4I-J). Supplementation with EB3 restored the Janusein-induced MT-loss phenotype in the EB1/2/3-knockout background (Figures 4I-J), supporting a functional cooperation between Janusein and EB proteins in regulating MT dynamics in vivo.

### Potential protofilament-binding interfaces within the Janusein NTD contribute to its MT-destabilizing activity

We next investigated the structural basis of Janusein’s NTD-driven MT destabilization by modeling its interaction with a short MT protofilament. AlphaFold3-based predictions suggested that the NTD consists of an α-helix flanked by two C2HC zinc-finger motifs (ZnF1 and ZnF2), both of which interact with β-tubulin (Figures 5A-B). Key residues in ZnF1 (R24, K33, K44, and R46) and ZnF2 (R127, R136, R147, and R151) are positioned near acidic tubulin surfaces, while the α-helix (R81, K83, R92, K95) interacts with α-tubulin via electrostatic contacts and a hydrophobic core (W80, H84, F87, I88, I91) (Figures 5C-F). These insights provided a framework for mutational testing.

**Figure 5.**
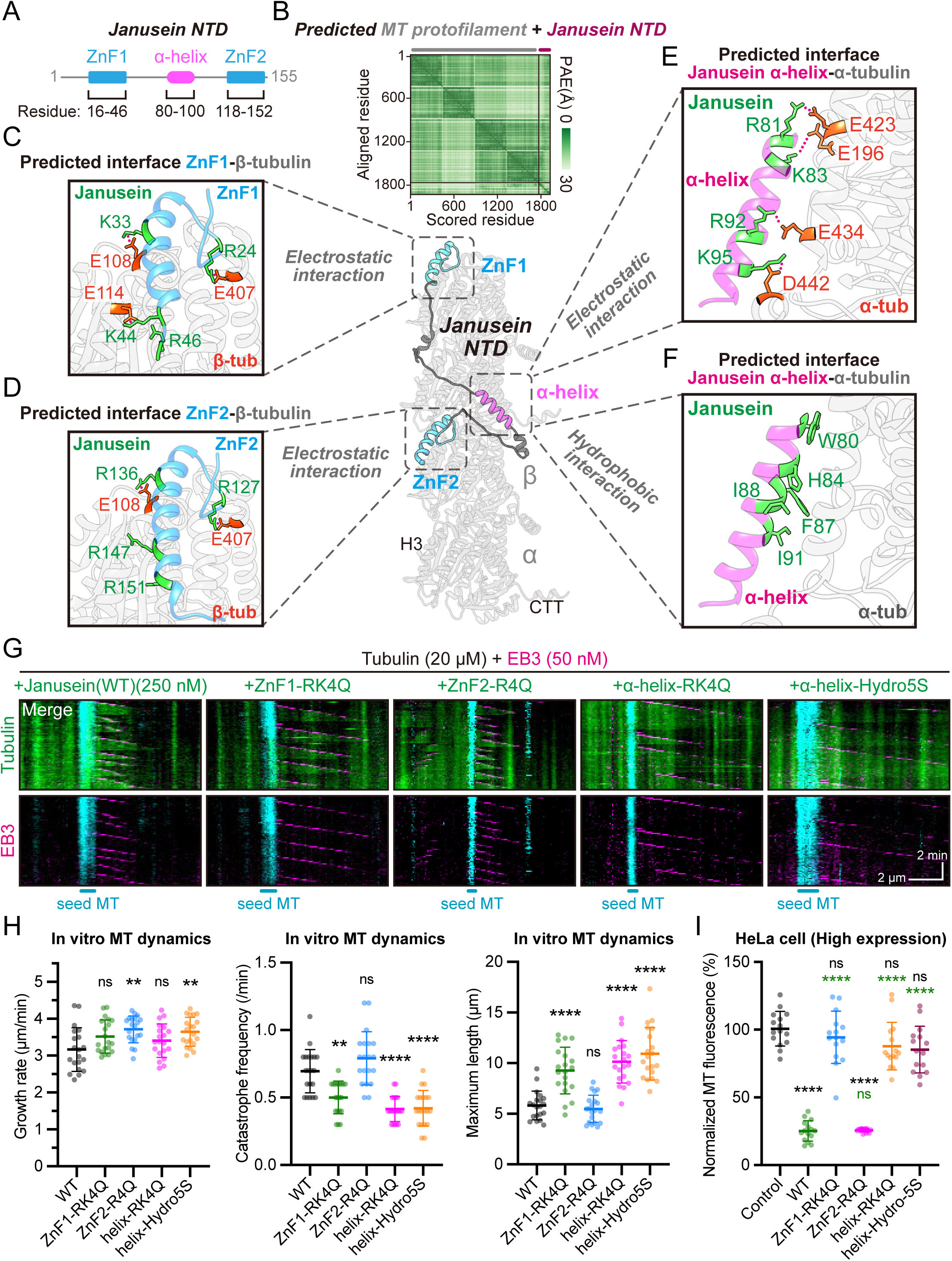
Interaction between the Janusein NTD and MT protofilament is critical for MT destabilization. (A) Schematic of the Janusein NTD (N-terminal domain), containing two zinc-finger motifs (ZnF1 and ZnF2, blue) and one α-helix (magenta). (B) AlphaFold3-predicted PAE (predicted alignment error) plot (top) and structure (bottom) of the MT protofilament and Janusein NTD. H3, helix 3; CTT, C-terminal tail. (C-D) Interaction pattern of the Janusein ZnF1 (C) or ZnF2 (D) with human β-tubulin. Key interacting residues are marked in green (Janusein) or orange red (tubulin). Salt bridges are indicated by dotted lines. (E-F) Electrostatic interaction (E) or hydrophobic interaction (F) pattern of the Janusein α-helix with human α-tubulin. Salt bridges are indicated by dotted lines. (G) Representative kymographs displaying in vitro MT dynamics with HiLyte647-conjugated seed MTs (cyan), HiLyte488-conjugated tubulins (green), EGFP-Janusein mutants (green) and EB3-mCherry (magenta). Scale bar, 2 μm (horizontal) and 2 min (vertical). (H) MT growth rate (left), catastrophe frequency (middle), and maximal growth length (right) for each group in (G) (N = 20). One-way ANOVA with Dunnett’s tests (left, right), or Kruskal-Wallis test with Dunn’s multiple comparisons (middle) were performed. (I) Normalized MT fluorescence in transfection-positive HeLa cells with high expression of WT Janusein, or Janusein mutants (ZnF1-RK4Q, ZnF2-R4Q, helix-RK4Q, helix-Hydro5S). Control represents untransfected cells in WT group. N = 15 cells. Shown as mean ± SD. Welch’s ANOVA with Dunnett’s tests were performed. ns, not significant; ** p < 0.01; **** p < 0.0001.

We generated mutants targeting predicted binding interfaces, including ZnF1-RK4Q (R24Q, K33Q, K44Q, R46Q), ZnF2-R4Q (R127Q, R136Q, R147Q, R151Q), α-helix-RK4Q (R81Q, K83Q, R92Q, K95Q), and α-helix-Hydro5S (W80S, H84S, F87S, I88S, I91S) (Figure S7A). In vitro reconstitution with EB3 revealed that mutations in ZnF1 or the α-helix reduced catastrophe frequency, while the ZnF2-R4Q mutant had minimal effect on MT catastrophe (Figures 5G-H). MT growth rates were also affected differently: ZnF2-R4Q and α-helix-Hydro5S mutants increased MT growth, while ZnF1-RK4Q and α-helix-RK4Q had subtle effects (Figure 5H). In HeLa cells, overexpression of ZnF1-RK4Q, α-helix-RK4Q, or α-helix-Hydro5S abolished the MT depolymerization observed with wild-type Janusein, while ZnF2-R4Q had minimal impact (Figures 5I and S7B). Charge-neutralizing single mutations within the α-helix produced only mild defects (Figure S7C), while single mutations within the hydrophobic core (W80S, H84S, F87S, I88S, or I91S) were sufficient to eliminate destabilizing activity (Figure S7D), highlighting the essential role of the α-helix’s hydrophobic interactions in Janusein’s MT destabilizing function.

Additionally, in *tba-5 (A19V); zchc-1A (null)* animals (*C. elegans*), expression of the corresponding ZCHC-1A (ZnF1-RK4Q) mutant failed to reverse the suppression of ciliary defects, whereas wild-type ZCHC-1A did (Figures S4F-G), consistent with impaired ZCHC-1A function in vivo. Together, these findings demonstrate that the ZnF1 and α-helix motifs are crucial for Janusein’s MT-destabilizing activity, whereas ZnF2 was dispensable for these activities under the conditions examined.

### A semi-flexible loop within the Janusein N-terminal domain contributes to lateral protofilament destabilization

Given that ZnF2 modulates, rather than drives, Janusein’s destabilizing activity, we next examined the Janusein (1-100) fragment, which retains ZnF1 and the α-helix but lacks ZnF2 (Figure 6A). This truncated version was sufficient to induce MT loss in HeLa cells (Figure 6B), demonstrating that the ZnF1-α-helix module alone can execute the core destabilizing function. To investigate the structural mechanisms, we performed all-atom molecular dynamics simulations using a two-protofilament MT system with and without Janusein (1-100) (Figure 6C; see Methods) (Fedorov et al., 2019; Xu et al., 2026a). In simulations, MT protofilaments maintained a stable conformation when simulated in isolation (Figure 6D). However, the addition of wild-type Janusein (1-100) led to a reduction in lateral contact stability between adjacent protofilaments (Figures 6D-E). These changes were absent when the MT-binding mutant ZnF1-RK4Q was introduced (Figures 6D-E; Figure S7E), suggesting that Janusein destabilizes lateral contacts through its NTD.

**Figure 6.**
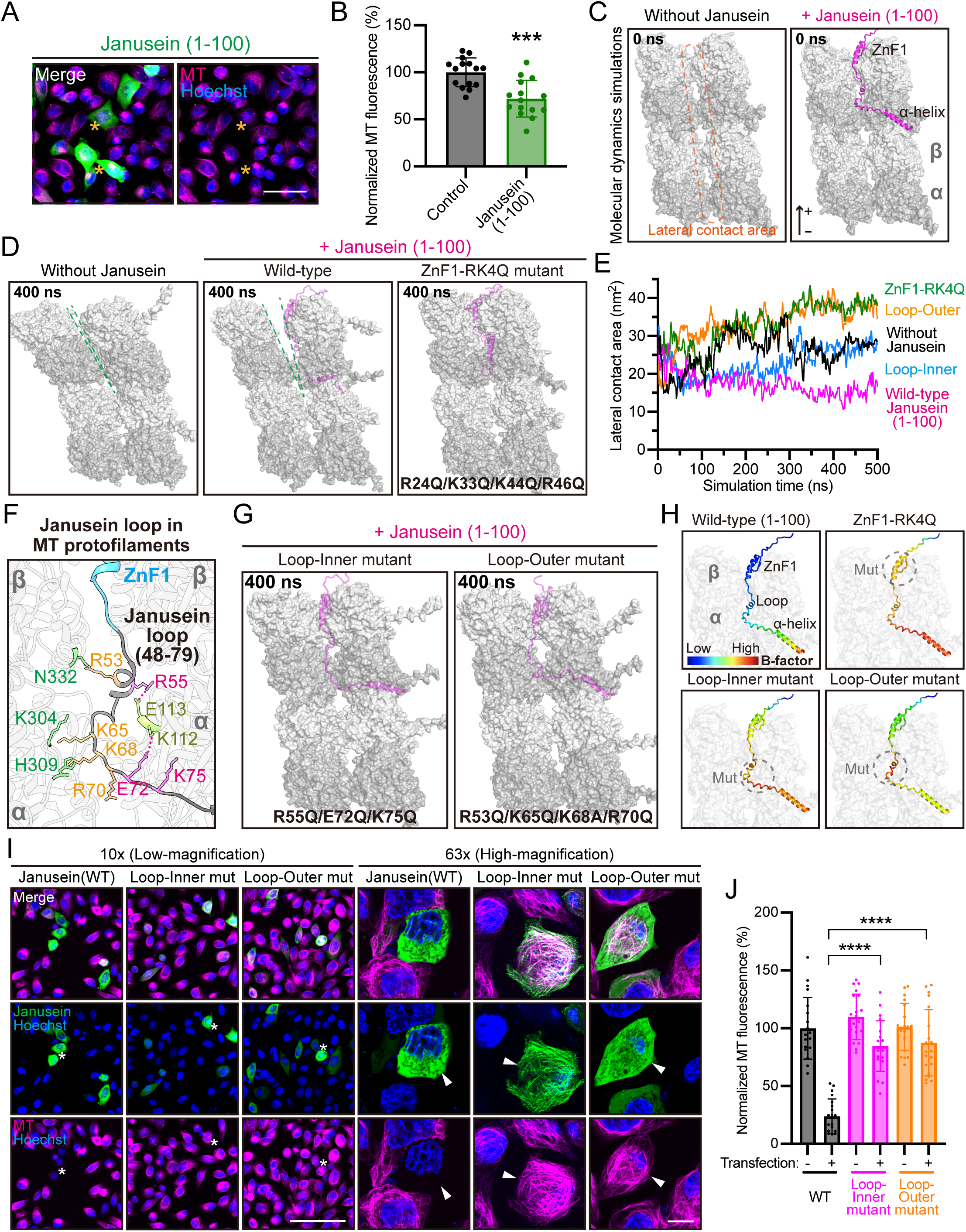
The semi-flexible loop of Janusein may disturb MT lateral contact. (A) Representative live-cell images of HeLa cells expressing EGFP-Janusein (1-100). Transfection-positive cells are indicated by orange asterisks. Scale bar, 50 μm. (B) Normalized MT fluorescence in control (untransfected cells) or Janusein (1-100)-expressing group. N = 15 cells. Shown as mean ± SD. Unpaired t tests were performed. *** p < 0.001. (C) Models of MT protofilaments without (left) or with (right) Janusein (1-100) (magenta) before molecular dynamics simulations. Each protofilament is composed of two α/β-tubulin heterodimers. Lateral contact area between two adjacent protofilaments is indicated by the orange box. +, MT plus end. (D) MT protofilaments without Janusein (left), or with wild-type Janusein (middle), or with the ZnF1-RK4Q mutant (right), after 400-ns simulations. The lateral surfaces of two protofilaments are marked by green dashed lines. (E) Statistics of lateral contact area of adjacent MT protofilaments in the absence or presence of Janusein mutants during simulations. (F) Interaction pattern between the Janusein semi-flexible loop (residues: 48-79) and tubulins. Key residues are marked in magenta (inward residues of Janusein), or orange (outward residues of Janusein), or yellow (interacting residues in tubulin), or green (repulsive residues in tubulin). Salt bridges are indicated by dotted lines. (G) MT protofilaments with the Janusein loop-inner mutant (left), or loop-outer mutant (right), after 400-ns simulations. (H) Distribution of residue-level B-factors along the Janusein (1-100) fragment. For each residue, the value is calculated relative to MTs during 500-ns simulations. For each system, the relative B-factors are normalized from blue (low, stable) to red (high, flexible). Mutated regions are indicated by gray circles. See also Movie S3. (I) Representative live-cell images of HeLa cells expressing wild-type Janusein, or loop-inner, or loop-outer mutants. For low-magnification fields (10x, left), transfection-positive cells are indicated by white asterisks. Scale bar, 100 μm. For high-magnification fields (63x, right), transfection-positive cells are indicated by white arrowheads. Scale bar, 10 μm. (J) Normalized MT fluorescence in each group in (I). The mean value of transfection-negative group of WT Janusein is normalized to 100%. N = 20 cells. Shown as mean ± SD. Unpaired t tests were performed. **** p < 0.0001.

Closer inspection of the Janusein MT-bound conformation revealed a semi-flexible loop adjacent to ZnF1 as a potential mediator of lateral destabilization (Figure 6F). The loop is predicted to have inward-facing residues (R55, E72, K75) that anchor the loop to the MT surface, while outward-facing residues (R53, K65, K68, R70) may disrupt interactions with the neighboring protofilament. Molecular dynamics simulations of loop mutants supported this model: the loop-inner mutant (R55Q/E72Q/K75Q) impaired loop anchoring, while the loop-outer mutant (R53Q/K65Q/K68A/R70Q) reduced steric clashes with the adjacent protofilament, both preserving lateral interactions in silico (Figures 6E and 6G; Figure S7E). Furthermore, we calculated relative B-factors to measure the conformational mobility of Janusein in the MT-bound state (Figure 6H; see Methods). Globally elevated B-factors in the ZnF1-RK4Q mutant were consistent with reduced MT binding, while the loop-inner mutant exhibited increased flexibility within the loop (Figure 6H; Movie S3). In contrast, the loop-outer mutant showed increased flexibility within the local mutation cluster (Figure 6H; Movie S3), confirming the functional relevance of the loop for Janusein-mediated MT destabilization.

We therefore tested this prediction in cells. Despite being expressed at levels comparable to those of wild-type Janusein, both the loop-inner and loop-outer mutants caused substantially less MT loss, with a greater proportion of the MT network retained in expressing cells (Figures 6I-J). Together, these cellular and simulation results implicate the semi-flexible N-terminal loop in Janusein-associated MT destabilization and support a model in which positioning of this loop at the lateral protofilament interface contributes to perturbation of lateral contacts.

### Mutations at the predicted Janusein-EB interface attenuate their functional cooperation

Given that EB3 potentiates Janusein’s MT-destabilizing activity in vitro (Figure 4E), we next investigated the molecular basis of this functional cooperation. AlphaFold3 modeling of full-length Janusein and EB3 predicted that the N-terminal calponin homology (CH) domain of EB3, instead of its C-terminal EB homology (EBH) domain, mediates the predominant interaction with the Janusein (1-100) fragment (Figure S7F). Therefore, we next focused on the interaction between EB3 CH domain and Janusein. AlphaFold3-based prediction involving EB3 CH domain and Janusein (1-100) suggested that they interact primarily through electrostatic contacts between basic residues in Janusein’s α-helix (R81, K83, R92, K95) and acidic residues in EB3 (D26, E13, D123, D120) (Figure 7A).

**Figure 7.**
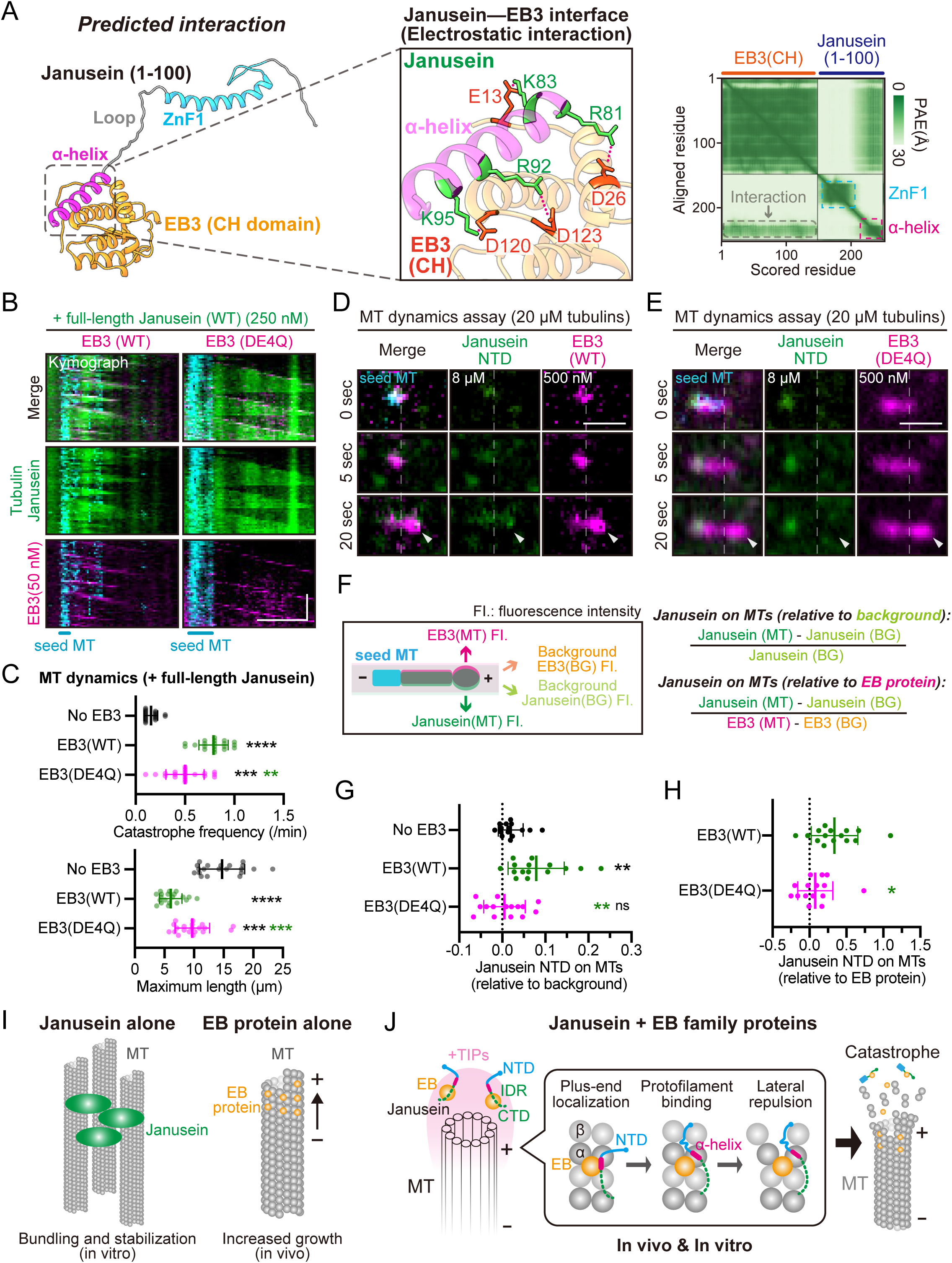
EB family proteins promote NTD accumulation at growing MT ends. (A) AlphaFold3-predicted structure (left) and PAE (predicted alignment error) plot (right) of the Janusein (1-100) fragment and EB3 CH (calponin homology) domain, highlighting the electrostatic interaction between the Janusein α-helix and EB3 CH domain. Salt bridges are indicated by dotted lines. (B) Representative kymographs showing in vitro MT dynamics with HiLyte488-conjugated tubulins (green) (20 μM), full-length EGFP-Janusein (green), and wild-type EB3-mCherry (magenta) (left), or EB3(DE4Q)-mCherry mutant (right). Scale bar, 2 μm (horizontal) and 2 min (vertical). (C) Quantification of MT catastrophe frequency (top) and maximal growth length (bottom) in each group in (B). N = 20. Shown as mean ± SD. Kruskal-Wallis test with Dunn’s multiple comparisons (top), or one-way ANOVA with Dunnett’s tests (bottom) were performed. (D-E) Representative images of in vitro MT dynamics assay (without methylcellulose) in the presence of EGFP-Janusein NTD and wild-type EB3 (D) or EB3 (DE4Q) mutants (E). HiLyte647-conjugated seed MTs and unlabeled tubulins (20 μM) were employed. White dashed lines mark the positions where dynamic MTs grow out. White arrowheads indicate the plus ends of dynamic MTs. Scale bar, 1 μm. (F) Schematic model demonstrating the quantification of Janusein on dynamic MT plus ends. Fluorescence intensities (FI.) of Janusein and EB3 on the MT plus ends, or in the background (BG) were measured. (G-H) Quantification of Janusein on MTs relative to background (G) or EB3 (H). N = 15 MTs. Shown as mean ± SD. Unpaired t tests were performed. ns, not significant; * p < 0.05; ** p < 0.01; *** p < 0.001; **** p < 0.0001. (I) Schematic model of Janusein alone (left) or EB protein alone (right) promoting MT growth. +, plus end. (J) Schematic model of the cooperative MT-destabilizing mechanism by Janusein and EB proteins. EB proteins interact with the Janusein α-helix (magenta) and promote the MT binding of Janusein NTD (N-terminal domain). Janusein inserts its semi-flexible loop into the interface between MT protofilaments, thus interfering with lateral contacts and triggers MT catastrophe.

To test this interaction model, we generated an EB3 mutant (DE4Q: E13Q/D26Q/D120Q/D123Q) designed to disrupt electrostatic interactions with Janusein’s α-helix (Figure S7A). Both wild-type EB3 and DE4Q mutant localized to growing MT ends in vitro (Figure 7B), confirming normal plus-end tracking. However, only wild-type EB3 robustly activated Janusein-driven catastrophe, while the DE4Q mutant showed a partial effect on catastrophe frequency [No EB3, 0.16 ± 0.07 min⁻¹; wild-type EB3, 0.79 ± 0.15 min⁻¹; EB3(DE4Q), 0.50 ± 0.19 min⁻¹] (Figures 7B-C). These results demonstrate that the Janusein-EB3 electrostatic interface is crucial for the functional cooperation between EB3 and Janusein in promoting MT catastrophe.

Given that the predicted EB-contacting interface lies within the Janusein NTD, we next asked whether the DE4Q substitutions in EB3 affected NTD accumulation at growing MT ends. Although Janusein NTD was sufficient to induce catastrophe in the presence of EB3 (Figures 4E-H), the isolated NTD showed little detectable enrichment on dynamic MTs even at 4 μM (Figures 4A-C). To improve signal detection, we therefore used 8 μM Janusein NTD and 500 nM EB3. Under these high-concentration imaging conditions, NTD enrichment at growing MT ends was lower with EB3 (DE4Q) than with wild-type EB3 (Figures 7D-H). Together, these results are consistent with a contribution of these acidic EB3 residues to NTD accumulation at growing MT ends and to functional cooperation between Janusein and EB3 in promoting MT catastrophe.

## Discussion

MT plus ends must sustain persistent growth while remaining poised for rapid destabilization. Our study identifies Janusein as a conserved factor that couples plus-end-associated growth to catastrophe through cooperative interactions with EB proteins. This synergy enables context-dependent tuning of MT behavior, allowing Janusein to function as a conditional destabilization factor depending on the local molecular environment.

The Janusein-EB module further expanded our conceptualization of MT depolymerization. Rather than being a purely stochastic event or simply the outcome of mechanical strain within the lattice, MT catastrophe emerges here as an actively licensed process that requires specific molecular assemblies at the plus end. Janusein and EB proteins together form a molecular switch that converts a growth-promoting environment into a destabilizing one. This reframes MT dynamics as a system governed not only by intrinsic polymer chemistry, but also by programmable protein logic operating at dynamic cellular interfaces.

Although EB proteins are demonstrated to increase MT catastrophe frequency in vitro (Bieling et al., 2007), they suppress MT catastrophe under cellular contexts and are traditionally viewed as growth promoters (Figure 7I) (Komarova et al., 2009). Our results suggest that EB proteins can license localized destabilization by recruiting and activating Janusein’s N-terminal module (Figure 7J). This cooperative logic allows catastrophe to be confined to regions of active growth, ensuring spatial precision and preventing indiscriminate MT loss. This study also reveals a dual role for EB proteins as both promoters of growth and gatekeepers of catastrophe, depending on the molecular partners they engage. Such duality exemplifies how the same plus-end platform can coordinate antagonistic activities with exquisite spatial precision.

Mechanistically, Janusein defines a mode of MT destabilization distinct from known pathways. Unlike severing enzymes (Hartman et al., 1998; McNally and Vale, 1993; Roll-Mecak and Vale, 2008; Vemu et al., 2018; Zehr et al., 2017), kinesin-13 motors (Desai et al., 1999; Friel and Howard, 2011; Wang et al., 2017), or stathmin (Belmont and Mitchison, 1996; Gupta et al., 2013), Janusein does not extract tubulin or peel protofilaments. Instead, modeling and mutational analysis suggest that Janusein perturbs lateral protofilament contacts through insertion of a semi-flexible loop. This structural interference lowers the energetic barrier for catastrophe without nucleotide hydrolysis, establishing ATP-independent catastrophe induction as a physiologically relevant principle.

The functional implications of this mechanism are particularly relevant for mitosis, where localized MT remodeling is essential for spindle assembly and integrity. Janusein depletion disrupts spindle organization, resembling phenotypes associated with loss of established depolymerases (Ganem and Compton, 2004; McNally et al., 2006), and suggesting that Janusein contributes to the finely balanced MT turnover required for mitotic fidelity. More broadly, EB-rich plus-end assemblies may serve as regulatory hubs where stabilizing and destabilizing forces are dynamically integrated rather than irreversible enzymatic remodeling.

Several limitations remain. The MT-bound conformation of Janusein has not yet been visualized directly, and the proposed lateral contact-interference mechanism is inferred from predictive modeling and simulations. High-resolution structural studies will be required to validate this model. In addition, Janusein likely operates within a broader regulatory network, and its activity may be modulated by additional cofactors or post-translational modifications.

In summary, Janusein represents an EB-dependent MT catastrophe factor that destabilizes MTs by locally perturbing protofilament interactions rather than by severing or peeling. Janusein exemplifies how cooperative interactions, rather than intrinsic enzymatic activity alone, can determine biological function. This highlights a broader principle in cell biology: emergent activities can arise from the assembly of components whose individual functions are non-destructive or even stabilizing. This idea is likely applicable beyond MTs, to actin dynamics, membrane remodeling, and signaling condensates.

## Materials and Methods

### Molecular biology

For mammalian transfection plasmids, cDNA construct of human *TUBA4A*, *TUBB8*, *MAPRE1 (EB1)*, *MAPRE3 (EB3)*, or *ZC2HC1A (Janusein)* was cloned into pCMV3.0 vector or pLV vector containing CMV enhancer, CMV promoter and bGH terminator, using In-Fusion® HD Cloning Kit (Takara Bio). Missense mutations or truncation mutants were introduced by linearizing this vector with 20 bp overlapped primers. The resulting PCR products were treated with *DpnI* digestion for 2 hours and then transformed into *E. coli Trans5α* (TransGen Biotech). The linearized PCR products were cyclized by spontaneous recombination in bacteria. Split-TurboID and full-length miniTurboID were adopted from previous studies (Addgene #116905, #153004, #153005) (Cho et al., 2020a). For microtubule (MT) inner proximity labeling system, split-TurboID N-terminal fragment (Turbo-N) was inserted into the G43/G44 residue pair of TUBA4A, flanked by ‘GSGSGA’ linkers on both sides. An HA tag (YPYDVPDYA) was inserted into the N-terminal side of Turbo-N. For MT outer proximity labeling system, Turbo-N was inserted into the C-terminus of TUBB8 with a ‘GGSGG’ linker. Split-TurboID C-terminal fragment (Turbo-C) was cloned into the pCMV3.0 empty vector, with C-terminal Flag (DYKDDDDK) tagged. Full-length miniTurboID was cloned into the pCMV3.0 empty vector, with N-terminal 3xFlag tagged. For purification of recombinant Janusein or EB3 from bacteria, cDNA of Flag-EGFP-Janusein-StrepII, or EB3-mCherry-StrepII was cloned into pET21 vector, downstream of T7 promoter (Addgene #89881). For purification of Janusein from SF9 insect cells, cDNA of Flag-EGFP-Janusein-StrepII was cloned into pFastBac vector, under the control of PH (polyhedrin) promoter and SV40 poly(A) terminator. For *C. elegans* overexpression plasmids, genome DNA construct of *zchc-1A*, along with 1,328-bp upstream promoter, was cloned into pPD95.77 vector containing *unc-54* 3’ UTR. GFP was fused to the C-terminus of ZCHC-1A and separated by a ‘GSGS’ linker. All plasmids were purified with AxyPrep Plasmid Purification Miniprep Kit (Axygen, #AP-MN-P-250) and PureLink Quick PCR purification Kit (Invitrogen, #K310001).

### Cell culture and transfection

HeLa and HEK293T cells were grown in DMEM medium (Yeasen) containing 10 % fetal bovine serum (FBS) (Gibco) and 1 % penicillin and streptomycin (Yeasen) at 37 ℃ with 5 % CO_2_. For imaging assays, cells were seeded on 35 mm glass-bottom confocal dishes (CellVis, #D35C4-20-0-N) the day before transfection. On the second day, cells with 50-70% confluence were transfected with *tuba4a (Turbo-N-i)* (300 ng), or *tubb8-Turbo-N* (300 ng), or *Turbo-C* (300 ng), or *tuba4a (Turbo-N-i)* (300 ng) + *Turbo-C* (300 ng), or *tubb8-Turbo-N* (300 ng) + *Turbo-C* (300 ng), or *miniTurboID* (150 ng), or *egfp-janusein* (400 ng), or *egfp-janusein* (400 ng) + *eb3-mCherry* (100 ng for low expression or in EB1/2/3 triple-knockout cells, or 400 ng for high expression), or *egfp-eb1* (400 ng) + *eb3-mCherry* (400 ng), respectively, using Lipofectamine^TM^ 3000 Transfection Reagent (Invitrogen) following manufacturer’s instruction. The dosages of mutants were consistent with their wild-type forms. For proximity labeling assays, cells were seeded on 100 mm cell-culture plates (Invitrogen, #150466) the day before transfection. On the second day, cells with 50-70% confluence were transfected with *tuba4a (Turbo-N-i)* (9 μg) + *Turbo-C* (9 μg), or *tubb8-Turbo-N* (9 μg) + *Turbo-C* (9 μg), or *miniTurboID* (4.5 μg) per plate, respectively, using PEI (polyethyleneimine) transfection reagent (Yeasen) following manufacturer’s instruction. To generate a stable HeLa cell line expressing EGFP-Janusein, pLV-EGFP-Janusein vector was co-transfected with psPAX2 and pMD2.G vectors into HEK293T cells at a molar ratio of 4:3:1. Lentivirus-containing medium was harvested and filtered through a 0.22-μm Millex GP filter unit. HeLa cells were subsequently infected with the lentivirus, and stable clones were selected by puromycin (1 μg/mL) treatment.

### Generation of the EB1/2/3 triple-knockout HeLa cell line

The EB1/2/3 triple-knockout HeLa cell line was a gift from Dr. Kai Jiang (Wuhan University). This cell line was generated using CRISPR-Cas9 technology through a sequential gene knockout strategy. First, the EB2 knockout HeLa cell line was constructed: HeLa cells grown in 6-well plates were transfected with 2 μg of PX459 carrying the single guide RNA (5’-CAGACATGACATCATTGCAT-3’) and selected with 8 μg/mL puromycin for 2 days. 5 days after the removal of puromycin, cells were diluted in 96-well plates to isolate single-cell colonies. Positive clones were screened by immunofluorescence and western blotting. Subsequently, using the EB2 knockout HeLa cell line as the parental cell line, EB1 (gRNA: 5’-AAGTCGACATGACATGCTGG-3’) and EB3 (gRNA: 5’-TGCACCTCAACTATACCAAG-3’) were individually knocked out in separate rounds, ultimately generating the EB1/2/3 triple knockout cell line.

### RNA interference (RNAi)

Short hairpin RNA (shRNA) sequences targeting Janusein were designed in BLOCK-iT™ RNAi Designer (Invitrogen). The 22-nt shRNA cassette was cloned into pLV vector under the control of U6 promoter, with inverted repeats separated by a ‘TCGA’ loop. A copGFP reporter was cloned into the same vector, downstream of the EF-1α (translation elongation factor) core promoter. HeLa cells were seeded on 35 mm glass-bottom confocal dishes (CellVis, #D35C4-20-0-N) the day before transfection. For each well, 400 ng of RNAi plasmid was transfected using Lipofectamine^TM^ 3000 Transfection Reagent (Invitrogen). After 8 h of incubation at 37 ℃ with 5 % CO_2_, nocodazole (Yeasen) was added into the culture medium at a final concentration of 50 ng/mL. After nocodazole treatment for 18 h, cells were washed 3 times with 1xPBS to remove drugs, and the medium was replaced with DMEM with 5% FBS. After an additional 40-min incubation at 37 ℃ with 5% CO_2_, cells were fixed with 4% paraformaldehyde (PFA) for subsequent immunofluorescence staining. The sequences of shRNA targets are: Janusein-shRNA#1: GCACATCCTCTAGTTAGTTTGC Janusein-shRNA#2: CATTCTTTCCAGTAGCATTAAC

### Genetic suppressor screen in *Caenorhabditis elegans*

*C. elegans* were cultured at 20 ℃ on standard nematode growth medium (NGM) plates with the *Escherichia coli* OP50. The wild-type (WT) strain was Bristol N2. The fluorescent dye DiI (1,1′-dioctadecyl-3,3,3′,3′-tetramethylindocarbocyanine perchlorate, Sigma) filling assay was widely used to assess the ciliary function (Inglis et al., 2007). *tba-5 (A19V)* mutant animals develop abnormal cilia and are dye-filling negative when cultured at 15 ℃ (Hao et al., 2011). Dye-filling assay was performed as previously described (Hao et al., 2011). We used forward genetic screening to isolate *tba-5 (A19V)* suppressors which exhibit rescued dye-filling phenotypes (dye-filling positive). Briefly, the mutant animals (P0) were synchronized at the L4 larval stage, collected in 4 mL M9 buffer, and incubated with 50 mM ethyl methanesulfonate (EMS) for 4 hours at room temperature with constant rotation. Then, animals were washed with M9 three times and cultured under standard conditions. After 24 hours, adult animals were bleached. Eggs (F1) were distributed and raised on about 1000 NGM plates, each containing 20-50 eggs. Adult F2 animals were maintained at 15 ℃ and subjected to dye filling. Homozygous dye-filling positive animals (suppressors) were then subjected to whole-genome sequencing (WGS) by Novogene Ltd. We identified *zchc-1A (Q78*)* as a candidate suppressor allele for the *tba-5 (A19V)* mutant.

### Transgenic overexpression in *Caenorhabditis elegans*

Extra-chromosomal transgenic lines of *C. elegans* were obtained by co-injecting P*zchc-1A::zchc-1A::gfp* (5 ng/μL, WT or ZnF1-RK4Q), P*dyf-1::osm-6::mScarlet* (10 ng/μL, cilia marker) and *rol-6 (su1006)* (20 ng/μL, selective marker) plasmids into the gonads of young adult worms. Homologous mutations of ZnF1-RK4Q in ZCHC-1A (the Janusein homolog) include R18Q (R24 for human Janusein), K27Q (K33 for human Janusein) and R40Q (R46 for human Janusein). At least 3 independent transgenic lines with a constant transmission rate (> 50%) were examined and analysed. To quantify amphid cilia length, *C. elegans* hermaphrodites were anesthetized with 0.1 mmol/L levamisole in M9 buffer, mounted on 3% agarose pads, and maintained at 15 ℃. Imaging was performed using an Axio Observer Z1 microscope (Carl Zeiss) equipped with 488 and 561 laser lines, a Yokogawa spinning disk head, an Andor iXon + EM-CCD camera, and a Zeiss 100×/1.46 objective. Images were acquired at 0.6-μm z-spacing by µManager (https://www.micro-manager.org). Images were taken using identical settings (EM-gain: 250, exposure time: 200 ms, 10 % of max laser). Image stacks were z-projected using the ‘max’ projection. Stacked images covering the whole cilia structures were used to measure the cilia length.

### Quantification of brood size

The *tba-5 (A19V); zchc-1A (Q78*)* double-mutant worms were crossed with wild-type (N2) animals to isolate homozygous *zchc-1A (Q78*)* single mutants. To minimize background mutations, the resulting *zchc-1A (Q78*)* mutants were backcrossed to N2 for three consecutive generations, with homozygous progeny selected at each generation. The resulting *zchc-1A (Q78*)* single mutants and their wild-type siblings were subjected for further analyses. For brood size analysis, L4-stage hermaphrodites were transferred singly to freshly seeded plates. Worms were repeatedly transferred to freshly seeded plates every 24 hours until reproduction ceased. We counted the number of hatched offspring at their L4 stages.

### Imaging

Super-resolution imaging was performed using Zeiss LSM900 with Airyscan2 confocal microscopy (Carl Zeiss) equipped with a 10×/0.45, or 63×/1.4 objective. Images were acquired at laser wavelength of 405 nm for Hoechst, 488 nm for EGFP/Dylight488, 561 nm for mCherry/Dylight549 and 640 nm for Deep Red/Dylight649. Images were taken using identical settings (Airyscan mode: Airyscan SR; Scan direction: bidirectional; Scan mode: frame). The cells were incubated with DMEM medium containing 2 μg/mL Hoechst 33342 and 1x Tubulin Tracker^TM^ Deep Red (Invitrogen, #T34076) at 37 °C for 30 min before live-cell imaging. For immunofluorescence (IF) staining, anti-DYKDDDDK first antibody (CST, #14793 or #8146) (1:800 diluted), anti-HA first antibody (CST, #2367) (1:100 diluted), anti-α-tubulin first antibody (Proteintech, #11224-1-AP) (1:200 diluted), anti-β-tubulin first antibody (Santa Cruz, #sc-58884, 1:200 diluted; or CST, #2146, 1:50 diluted), were used. After incubation with first antibodies overnight at 4 ℃, cells were washed three times with PBS buffer. Secondary antibodies (Dylight488/549/649, Abbkine, 1:250 diluted) and Dylight488-conjugated Streptavidin (Invitrogen, #21832, 1:500 diluted) were added for 1-hour incubation at RT, followed by washing three times with PBS. The cells were incubated with 1 μg/mL Hoechst 33342 for 10 min before imaging. To measure Janusein and EB3 velocities in HeLa cells, only cells with low expression of EB3 (exhibiting clear plus-end tracks) were selected. Imaging was performed using an Axio Observer Z1 microscope (Carl Zeiss) equipped with 488 and 561 laser lines, a Yokogawa spinning disk head, an Andor iXon + EM-CCD camera, and a Zeiss 100×/1.46 objective. Images were acquired at 1-sec intervals by µManager (https://www.micro-manager.org). Images were taken using identical settings (EM-gain: 250, exposure time: 250 ms, 10 % of max laser). Image analysis and measurement were performed with Fiji / ImageJ software (http://rsbweb.nih.gov/ij/). The visualized color of red channel was changed to colorblind-safe magenta in ImageJ. All images shown were adjusted linearly. Kymographs were generated in Fiji by manually drawing lines along EB3 traces.

### Colchicine treatment

HeLa cells seeded on 35 mm glass-bottom confocal dishes (CellVis, #D35C4-20-0-N) were transfected with either EGFP (400 ng) or EGFP-Janusein (△NTD) (400 ng). After 18 hours, cell-culture medium was replaced with DMEM medium (+ FBS) containing either 5 μM colchicine (HEOWNS, #C-93541) or DMSO (control group), followed by incubation for 30 min at 37 ℃ with 5 % CO_2_. Next, cells were washed with 1xPBS for 3 times, and fixed with 4% PFA for subsequent immunofluorescence staining.

### Purification of recombinant proteins

For expression and purification of Flag-EGFP-Janusein-StrepII and EB3-mCherry-StrepII from bacteria, plasmids were transformed into BL21 (DE3) competent cells (TransGen Biotech). 10-mL culture from a single colony was used to inoculate a 1-L fresh LB medium for further incubation at 37 °C with shaking. After 3.5-4 h, IPTG was added to the culture to a final concentration of 1 mM to induce protein expression at 16 °C for 16 h. For purification of Flag-EGFP-Janusein-StrepII from Sf9 insect cells, baculovirus were constructed using Bac-to-Bac® baculovirus expression system (Invitrogen) following manufacturer’s instruction. 700 mL of infected Sf9 cells in Sf-900™ II SFM medium (Thermo Fisher Scientific) were employed for purification.

For Janusein, bacteria or insect cell pellets were collected by centrifugation and resuspended in 50 mL of ice-cold lysis buffer (20 mM HEPES, 500 mM NaCl, 5 % glycerol, pH 8.0) containing 1 mM PMSF. For EB3, the lysis buffer was BRB80 (80 mM PIPES, 100 mM KOH, 1 mM EGTA, 1mM MgCl_2_, pH 6.9) containing 500 mM NaCl and 1 mM PMSF. After breaking cells by microfluidizer (AH-TITAN PLUS) at 800 bar for 15 min, the lysate was centrifuged at 10,000 rpm for 40 min at 4 °C. The supernatant was transferred to a fresh tube containing 4 mL of 50 % slurry of Streptactin beads 4FF (Smart-Lifesciences). After rotating at 4 °C for 2 h, the mixture was transferred to a column to allow the settlement of the beads. The column was washed with 50 mL lysis buffer, and eluted with 10 mL elution buffer (lysis buffer with 2.5 mM dethiolated biotin). The Janusein-containing eluates were then incubated with 500 μL Anti-DYKDDDDK affinity beads (Smart-Lifesciences) at 4 °C for 1 h, washed with 10 mL lysis buffer, and eluted with 4 mL elution buffer (lysis buffer with 0.25 mg/mL 3xFlag peptide). Final eluates were concentrated to 50-100 μL with Amicon Ultra 30K device (Millipore) at 8 °C. During this step, dethiolated biotin or 3xFlag peptide was removed by buffer exchange. Protein concentrations were determined by Bradford assay. The proteins were divided into 5 μL aliquots, snap-frozen in liquid nitrogen and stored at -80 °C.

### In vitro microtubule binding

GMPCPP-stabilized MT polymerization followed the protocol described previously (Chen et al., 2025). Briefly, MRB80 buffer (BRB80 buffer with 4 mM MgCl_2_) containing 8 μM rhodamine-labeled porcine brain tubulins and 1 mM GMPCPP (JenaBiosciences) were incubated at 37 ℃ for 30 min. After centrifugation at 15,800 rpm for 30 min, the pellet (containing polymerized MTs) was suspended with MRB80 buffer and then flowed into anti-β-tubulin (Santa Cruz, #sc-58884) enveloped chambers for immobilization. The surfaces were then blocked by MRB80 buffer containing 1% Pluronic F-127 (Invitrogen). Next, HEPES buffer containing 150 mM NaCl, 1 mM ATP (Macklin) and 6 μM EGFP-Janusein was directly flowed into the chamber, and then analysed by a confocal microscope.

### In vitro microtubule dynamics

MT dynamics assay followed the protocol described previously (Zhang et al., 2022). In brief, the flow chamber was incubated first with 0.2 mg/mL PLL-PEG-biotin (Susos AG) and then with 1 mg/mL neutravidin (Invitrogen) in MRB80 buffer. Subsequently, rhodamine (or HiLyte488, or HiLyte647)-conjugated and biotin-conjugated (Cytoskeleton, Inc) MT seeds were attached to the coverslip through biotin–neutravidin interactions. The flow chamber was further blocked with 1 mg/mL κ-casein. The reaction mixture, which consisted of MRB80 buffer containing EGFP-Janusein (or its mutant), EB3-mCherry (or its mutant), porcine brain tubulins (20 μM, unless otherwise noted) (Cytoskeleton, Inc), 2 mM GTP, 0.2 mg/ml κ-casein, 0.1% methylcellulose (in Figure 7D-E, methylcellulose was not used), and oxygen scavenger mix (50 mM glucose, 400 μg/ml glucose oxidase, 200 μg/ml catalase, and 4 mM DTT) was added to the chamber. For fluorescent tubulins, the percentage of HiLyte488-conjugated tubulins (Cytoskeleton, Inc) is 12.5%, and the percentage of rhodamine-conjugated tubulins (Cytoskeleton, Inc) is 2.5%. The flow chamber was sealed with vacuum grease and imaged immediately at 30 °C using a Nikon Eclipse Ti-E inverted microscopy equipped with a 100×/1.4 objective. Images were acquired at the TIRF (total internal reflection fluorescence) mode with exposure time of 200 ms, reflection angle of 65°, frame interval of 8 sec, and laser wavelength of 488 nm / 561 nm / 647 nm. All experiments were independently repeated three times with consistent results.

### Proximity labeling and mass spectrometry

Two 100-mm cell-culture plates of cells (about 10^7^ cells) (HeLa or HEK293T) were employed in one replicate. Each group (negative-control, inner or outer) encompassed two independent replicates. After transfection of proximity labeling-associated plasmids for 18 hours, D-biotin was added into the DMEM culturing medium to a final concentration of 50 μM, followed by incubation at 37 ℃ and 5 % CO_2_ for 3 hours. Then, cells were washed with ice-cold PBS buffer for 3 times to stop the reaction and lysed in lysis buffer (Beyotime, #P0013) with 1% Triton X-100 and 1mM PMSF. Four volumes of 1xPBS buffer with 5% glycerol and 1mM PMSF were gently mixed with the lysates, which contained the biotinylated proteome. The lysates were centrifuged for 10 min at 12,000 rpm at 4 °C. After centrifugation, lysate supernatants were mixed with pre-washed streptactin beads 4FF (SmartLifesciences; 200 μL bed volume per sample), and incubated at 4 °C on a shaker for 2 h. After incubation, samples were transferred to a column to allow the settlement of the beads. The beads were washed with 10 mL 1xPBS buffer, and then eluted with 4 mL 1xPBS buffer containing 2.5 mM D-biotin. Final eluates were concentrated to 50 μL with Amicon Ultra 3K device (Millipore) at 4 °C. During this step, D-biotin was removed by buffer exchange without losing the majority of proteins.

For liquid chromatography-tandem mass spectrometry analysis, trypsin-digested peptides were separated by an EASY-nLC system (Thermo Fisher Scientific) coupled to a Q Exactive mass spectrometer (Thermo Fisher Scientific). The analytical column was a fused-silica capillary column (75 μm in internal diameter, 15 cm length; Upchurch) packed with C-18 resin (300 A°, 5 mm; Varian). The Q Exactive mass spectrometer was operated in data-dependent acquisition mode under Xcalibur control, and there was a single full-scan mass spectrum in the orbitrap (400-1,800 m/z, 60,000 resolution) followed by the top 10 most intense precursor ions subjected to HCD fragmentation at 27% normalized collision energy. The tandem mass spectrometry spectra were searched against the UniProt human proteome database (2024_06 release) using Proteome Discoverer (Thermo Fisher Scientific). Peptide Spectrum Matches (PSMs) were validated using a maximum false discovery rate (FDR) threshold of 1.0%.

### Data processing and differential abundance analysis

For each sample (replicate), the total PSMs (peptide spectrum matches) were normalized to NC (negative control) group (Figure S2E). Normalized PSMs were subjected to differential abundance analysis based on DEseq2 R package. Candidate MT-associated proteins (MAPs) were defined using the following cutoffs: BaseMean (average PSM per protein) > 1.0, log_2_FoldChange > 0.7, P value < 0.5 (different from the thresholds used in RNA sequencing). Principal component analysis (PCA) was performed by the ‘plotPCA’ function in DEseq2 R package. Venn diagrams were generated by the VennDiagram R package.

### Western blot

Eluates from streptactin beads (the purified biotinylated proteome) and total lysates from HeLa cells were analysed by SDS-PAGE followed by Western blotting. First antibodies included anti-β-actin (Proteintech, #20536-1-AP; 1:5,000), anti-EML4 (Proteintech, #22709-1-AP; 1:1,000), anti-JPT1 (Proteintech, #14914-1-AP; 1:500), anti-Janusein (CGI-62) (Invitrogen, #PA531854; 1:1,000). After incubation with first antibodies overnight at 4 ℃ on a shaker, the PVDF membranes were washed three times with 1xTBST buffer (10 min per wash on a shaker). Secondary antibodies were added at dilutions of 1:10,000 (goat anti-rabbit/-mouse, EasyBio), or 1:2,000 (HRP-conjugated streptavidin, Proteintech) and incubated for 1 h at room temperature on a shaker, followed by 3 washes with 1xTBST buffer. Finally, chemiluminescence (SuperSignal™ West Pico PLUS, Thermo Fisher Scientific) agent was added and signals were visualized using a FluorChem illuminator (ProteinSimple).

### AlphaFold3-based structural prediction and pDockQ calculation

AlphaFold3 was utilized to predict interactions between MTs and candidate MAPs (<u>alphafoldserver.com/</u>) (Abramson et al., 2024). Protein sequences were obtained from the Uniprot database (https://www.uniprot.org/). For each prediction, two human α-tubulins (TUBA4A), two human β-tubulins (TUBB8), as well as one candidate MAP, were submitted as multimers to AlphaFold3. The predicted multimer structures and predicted alignment error (PAE) plots were subsequently analysed. Interaction strength was defined based on the average PAE value of the putative binding region: candidates were classified as ‘*Weak binding*’ if the region exhibited potential contacts with the MT model (comprising two tubulin heterodimers) but had an average PAE of 15-25 Å; ‘*Strong binding*’ if the average PAE was < 15 Å; and ‘*No binding*’ if the average PAE exceeded 25 Å. Interaction reliability was further assessed using the predicted DockQ (pDockQ) score, a quantitative measure of the confidence of AlphaFold-predicted protein-protein interactions (Bryant et al., 2022). The MT model (two TUBA4A and two TUBB8) was designated as a monomer, and pDockQ score was calculated for the putative ‘MT’-MAP dimer.

The interaction between Janusein (ZC2HC1A) and MT protofilament (two tubulin heterodimers) was predicted by AlphaFold3 using 4 additional magnesium ions (Mg^2+^), 4 additional GTP molecules and 2 additional zinc ions (Zn^2+^) as input. The interactions between Janusein and other plus-end-binding proteins such as EB3 were predicted as multimer models.

### Multiple sequence alignment (MSA)

Janusein protein sequences from various model organisms were retrieved from the National Center for Biotechnology Information (NCBI) (www.ncbi.nlm.nih.gov/). Human Janusein (ZC2HC1A) homologs were identified using the NCBI protein BLAST tool. For each organism, the top-scoring candidate homologs were selected and subsequently subjected to a reciprocal BLAST against the human proteome to confirm the orthology. Only candidates whose highest-ranking human homolog was Janusein (ZC2HC1A) were considered bona fide Janusein homologs. The validated Janusein homologs include: ZC2HC1A (Uniprot ID: Q96GY0) (human; *Homo sapiens*), Zc2hc1a (Q8BJH1) (mouse; *Mus musculus*), ZCHC-1A (Q22122) (worm; *Caenorhabditis elegans*), Zc2hc1a (Q5PPV5) (frog; *Xenopus laevis*), Zc2hc1a (Q7SXT7) (zebrafish; *Danio rerio*), Zc2hc1a (A0A8V0XDQ2) (chicken; *Gallus gallus*), Zc2hc1a (A0A7M7N3H1) (sea urchin; *Strongylocentrotus purpuratus*), CG31554 (Q9VNI7) (fly; *Drosophila melanogaster*), At2g02148 (Q56XX3) (*Arabidopsis thaliana*), 04g217952v5 (A0A2K3DTI5) (algae, *Chlamydomonas reinhardtii*). Sequence alignment of these homologs was performed using Clustal X2.1 (http://www.clustal.org/), and a phylogenetic tree was generated in Jalview 2.11.2.1.

### Identification of critical interaction residues

UCSF ChimeraX (https://www.rbvi.ucsf.edu/chimerax/) was employed to identify critical residues mediating Janusein-MT and Janusein-EB3 interactions. Residue pairs from different chains were considered candidate interaction sites if their side-chain atoms exhibited van der Waals (VDW) overlap < 0.6 Å. The credibility of these interactions was further assessed based on residue conservatism, side-chain orientation, charge and hydrophilism characteristics. Conserved residues with high interaction probability were then selected. These included R24, K33, K44, R46 (Janusein ZnF1); R127, R136, R147, R151 (Janusein ZnF2); R81, K83, R92, K95 (Janusein α-helix); K313, E317, R321 (Janusein CTD); R55, E72, K75 (Janusein semi-flexible loop); E13, D26, D120, D123 (EB3 CH domain). Moreover, W80, H84, F87, I88 and I91 within the Janusein α-helix are highly conserved across species, and identified as critical residues forming a hydrophobic core essential for Janusein-MT and Janusein-EB3 interactions. R53, K65, and K68/R70 within the Janusein semi-flexible loop have potent steric/electrostatic clashes with N332 (β-tubulin), K304 (α-tubulin) and H309 (α-tubulin), respectively. Structural representations were generated using UCSF ChimeraX.

### Structural modeling of Janusein-MT for molecular dynamics simulations

The structure of Janusein binding to a protofilament consisting of two TUBA4A and TUBB8 heterodimers was predicted by AlphaFold3. Using the GMPCPP-stabilized MT structure (PDB: 3J6E) as template, we aligned the predicted Janusein-bound protofilament to the tubulin wall resolved by Cryo-EM. We performed homology modeling through the SWISS-MODEL workspace (Waterhouse et al., 2018) to ensure the tubulins we used (TUBA4A and TUBB8) to fit the conformation of solved structure. To simulate the physiological GTP-bound conditions, we modified the GMPCPP ligand in PyMOL v2.5 (http://www.pymol.org/) by replacing the methylene group bridging the α-and β-phosphates with an oxygen atom. This conversion was achieved using PyMOL’s molecular builder toolkit with geometry optimization to maintain proper bond lengths and tetrahedral coordination angles. Ionizable residue protonation states were subsequently determined using PDBFixer (http://github.com/openmm/pdbfixer). All structural manipulations preserved the original MT lattice parameters observed in the template crystal structure.

### Molecular dynamics simulations

A virtual three-dimensional cubic reaction volume filled with TIP3P water and periodic boundary conditions was used for all the simulations. The box dimensions were set such that the distance from any protein surface to the nearest box edge was at least 2 nanometers, ensuring sufficient solvation. To mimic physiological conditions, the ionic strength of the solution was adjusted to 100 mM by supplementing K⁺ and Cl⁻ ions, and the total charge of the system was neutralized. Simulations were performed using the GROMACS 2024.2 software package (http://www.gromacs.org/) with the CHARMM36m force field (Huang et al., 2017). Parameters for GTP were generated using the CHARMM General Force Field (CGenFF) via the CHARMM-GUI platform (http://www.charmm-gui.org/).

Each simulation system was prepared as described in the ‘Structural modeling of Janusein-MT for molecular dynamics simulations’ section, followed by energy minimization using the steepest descent algorithm. Subsequently, a two-step equilibration was performed: (1) a 100 ps simulation under NVT conditions using the V-rescale thermostat (τ = 0.1 ps, T = 300 K); and (2) a 100 ps simulation under NPT conditions using the same thermostat and the C-rescale barostat (τ = 2.0 ps, P = 1 bar). During both equilibration steps, position restraints were applied to the heavy atoms of the protein and ligand. Production simulations were performed in the NPT ensemble at 300 K for 1,000 ns, using the V-rescale thermostat and C-rescale barostat. Positional restraints were applied to the Cα atoms of TUBA4A chains at the minus end of the tubulin complex throughout the production runs. Long-range electrostatic interactions were treated with the Particle Mesh Ewald method. All bonds involving hydrogen atoms were constrained using the LINCS algorithm, allowing a 2-fs integration time step. Mutations in each Janusein chain were introduced in PyMOL v2.5.

### Lateral contact area calculation

The total contact area between two simulated protofilaments was estimated based on their Solvent Accessible Surface Areas (SASA). For the control group which does not contain a Janusein, the contact area (A) was calculated as:

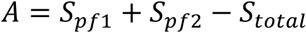

Where *S_pf_*_1_ and *S_pf_*_2_ are the SASA values of the individual protofilaments, and *S_total_* is the SASA of the protofilament complex.

Because of the position Janusein located, which is partly the lateral interface between protofilaments, the contact area was calculated as:

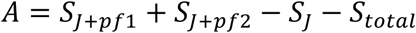

Where *S_J_*_+*pf*1_ and *S_J_*_+*pf*2_ are the SASA values of the protofilament-Janusein complexes, and *S_J_* is the SASA values of the Janusein. SASA values were computed using the GROMACS sasa module with default probe radius (0.14 nm) and parameters.

### B-factor calculation

Atom-level B-factors were calculated as:

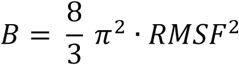

Where RMSF (Root Mean Square Fluctuation) values were obtained with the GROMACS rmsf module. Residue-level B-factors were then derived by averaging atom-level B-factors across all atoms within each residue.

### Quantification and statistical analysis

Sample sizes were determined based on prior published studies of similar design. GraphPad Prism 9 (GraphPad Software, Inc.) was used for statistical analyses. For comparisons between two groups (Gaussian distribution), unpaired t tests were employed. For datasets lacking Gaussian distribution, nonparametric Mann-Whitney U tests were employed. For comparisons between multiple (>2) groups with Gaussian distribution and equivalent standard deviation (SD), one-way ANOVA with Dunnett’s (compared with one control group) or Tukey’s (compared with each other) multiple comparisons tests were performed. For multiple (>2) groups with Gaussian distribution but inequivalent SD, Welch’s ANOVA analyses with Dunnett’s or Tukey’s multiple comparisons tests were performed. For multiple (>2) groups lacking Gaussian distribution, nonparametric tests (Kruskal-Wallis test with Dunn’s multiple comparisons) were performed. N represents the number of samples per group. Statistical significances are designated as: ns p > 0.05, * p < 0.05, ** p < 0.01, *** p < 0.001 and **** p < 0.0001.

## Acknowledgements

This work was supported by the National Key R&D Program of China (2024YFA1307301, 2022YFA1302700) and the National Natural Science Foundation of China (92254306 to G.O.; 32600008 and 323B200173 to K.X.; 32270773 and 32470730 to W.L.; 32430026, 32021002, and 31991191 to G.O.). We thank Dr. Xin Liang (Tsinghua University) for his assistance with microtubule dynamics assays. We appreciate Dr. Xuebiao Yao (USTC), Dr. Andrew Carter (MRC-LMB), Dr. Carsten Janke (Institut Curie) for their insightful discussions on this project. We are grateful to the Protein Chemistry and Proteomics Facility at Tsinghua University (particularly Xianbin Meng, Xiaolin Tian, and Meng Han), and the Facility of Cell Imaging at the Tsinghua University (especially Bingyu Liu and Jinyu Wang), for support with proteomic sample processing and analysis, super-resolution imaging, and TIRF imaging.

## Author Contributions

K.X. and G.O. conceived and designed this study. G.O. supervised this study. K.X. and Y.C. performed mass spectrometry-based profiling. Z.G. performed molecular dynamics simulations. K.X. performed other experiments. Y.Y. assisted in live-cell imaging. C.G. and K.J. provided the EB1/2/3 knockout cell line. K.X. analysed the data. K.X., W.L. and G.O. wrote this manuscript.

## Declaration of Interests

The authors declare no competing interests.

## Data Availability

All data in this study are available upon reasonable request.

---

**Figure S1.**
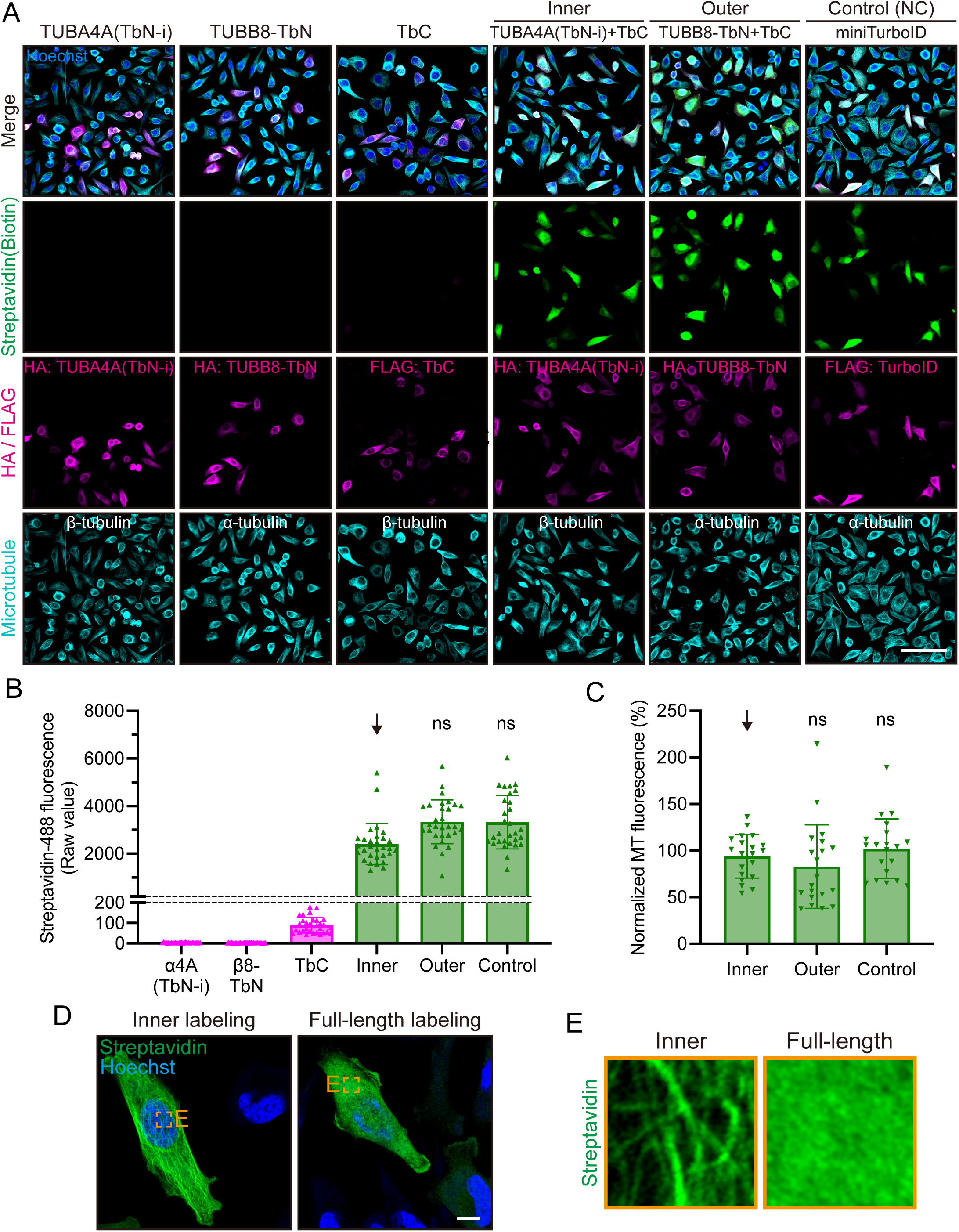
Split-TurboID proximity labeling systems have minimal impact on cellular MT networks. (A) Representative immunofluorescence images of different proximity labeling systems in HeLa cells. Engineered constructs (shown in Figure 1B) are directly transfected into HeLa cells for over-expression. Streptavidin (green) marks the biotinylated proteome in cells. HA or Flag antibodies label the engineered TurboID constructs. Hoechst (blue) marks the nucleus. Scale bar, 100 μm. (B) Dylight 488-conjugated streptavidin fluorescence (raw value) in each group in (A). N = 30 transfection-positive (Anti-HA or -Flag staining positive) cells. Shown as mean ± SD. Unpaired t tests were performed. (C) MT fluorescence in inner, outer or NC group. For each individual cell, the whole-cell MT fluorescence (Anti-α-tubulin or -β-tubulin staining) is measured. For each group, the average MT fluorescence of 20 randomly selected transfection-negative cells is normalized to 100%, and unpaired t test is performed to compare the statistical difference between transfection-negative cells (not shown, N = 20) and transfection-positive cells (shown in the panel, N = 20). Shown as mean ± SD. Unpaired t tests were performed. ns, not significant. (D) Representative immunofluorescence images showing Streptavidin-stained biotinylated proteome in inner or full-length labeling group. Scale bar, 10 μm. (E) Enlarged view of Streptavidin staining in (D).

**Figure S2.**
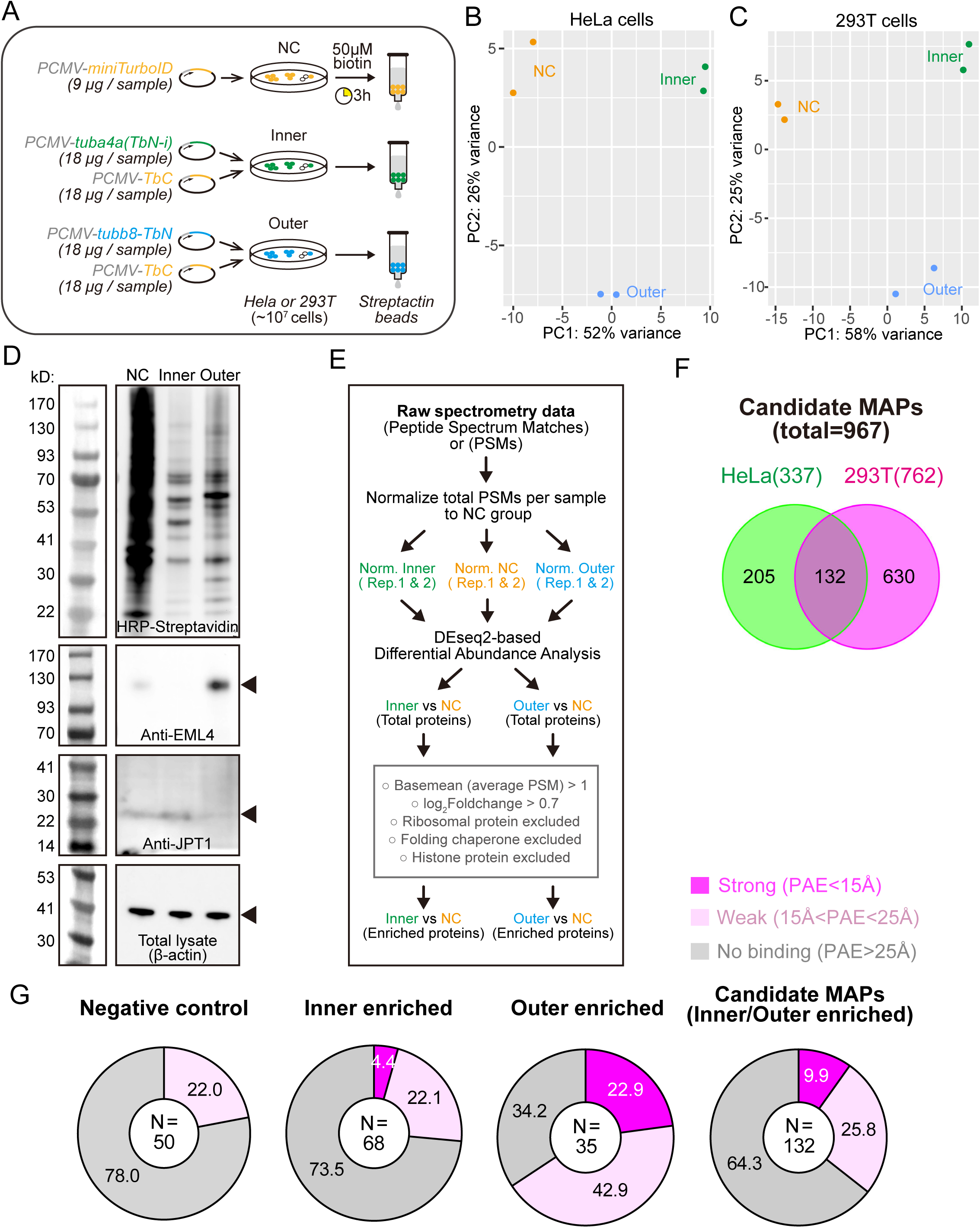
Workflow for mass spectrometry-based profiling of MAPs. (A) Schematic of the workflow for proximity labeling and affinity purification of the biotinylated proteome from HeLa or HEK293T cells (see Methods). Transfection doses are adopted from immunofluorescence assays in Figure S1A, and scaled up to accommodate 10^7^ cells. (B-C) Principal component analysis (PCA) plots showing clear separation between NC, inner and outer groups (two independent technical replicates for each group) for HeLa (B) or HEK293T (C) cells. (D) Western blot analysis of the biotinylated proteome (HRP-streptavidin), MT-outer protein EML4 (Anti-EML4), and MT-inner protein JPT1 (Anti-JPT1) in different labeling groups after proximity labeling and affinity purification from HeLa cells, as demonstrated in (A). Target proteins (EML4, JPT1 or β-actin) are indicated by black arrowheads. (E) The pipeline for DEseq2-based differential abundance analysis to identify inner/outer-enriched candidate MAPs (see also Methods). Briefly, total peptide spectrum matches (PSMs) for each replicate in inner and outer groups were first normalized to the corresponding NC group, and then subjected to DEseq2-based differential expression (abundance) analysis. Proteins passing empirical thresholds (average PSM>1, log2FC>0.7, excluding ribosomal proteins, folding chaperones, histones) were filtered out as candidate MAPs. (F) Venn diagram indicating the number of all candidate MAPs enriched in HeLa cells and HEK293T cells, as well as those shared in both cell lines (N = 132). (G) Pie charts showing the percentages of candidate MAPs classified as ‘Strong’ (magenta), ‘Weak’ (pink), or ‘No binding’ (gray) to MTs, according to the average PAE (predicted alignment error) values for the putative MT-MAP interaction regions. A total of 132 MAPs identified in both HeLa and HEK293T cells were subjected to AlphaFold3-based prediction.

**Figure S3.**
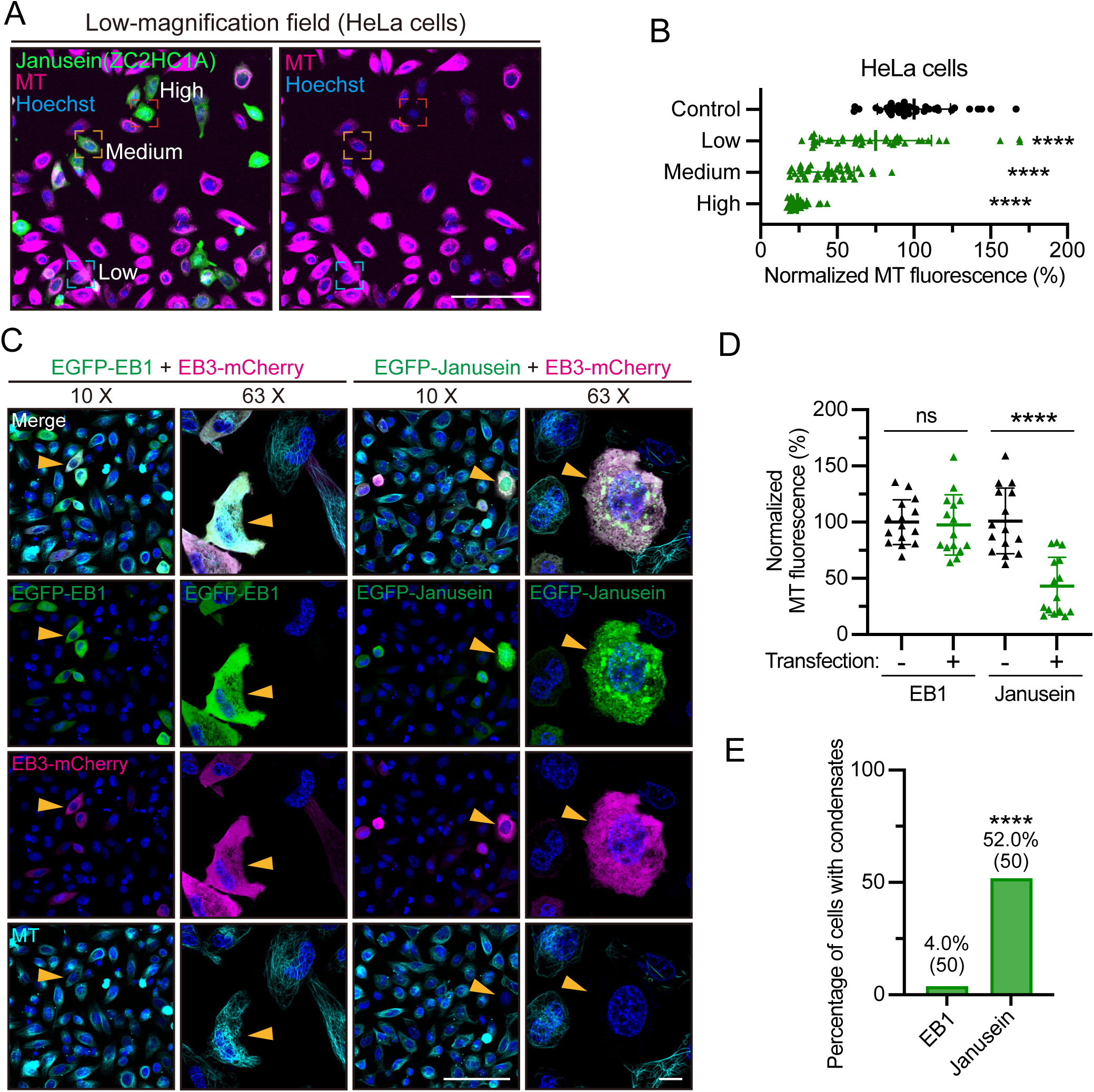
Overexpression of Janusein destabilizes MT networks in HeLa cells. (A) Low-magnification overview of EGFP-Janusein-transfected HeLa cells. Tubulin Tracker Deep Red (magenta) is used to stain MT networks. Representative cells with low (blue), medium (orange), or high (red) Janusein expression are indicated by differentially colored boxes. See also Figures 2H-2I for the quantitative classification of low, medium, or high Janusein expression. Scale bar, 100 μm. (B) Whole-cell MT fluorescence intensities in control (transfection-negative cells), low-, medium-, or high-Janusein expression cells. The average value of control group is normalized to 100%. Shown as mean ± SD. N = 40 cells. One-way ANOVA with Dunnett’s tests were performed. (C) Representative live-cell images of HeLa cells over-expressing EGFP-EB1 and EB3-mCherry (left), or EGFP-Janusein and EB3-mCherry (right). Transfection-positive cells are indicated by orange arrowheads. Tubulin Tracker Deep Red is used to stain MT networks. Scale bar, 100 μm (10x) or 10 μm (63x). (D) Whole-cell MT fluorescence intensity in each group in (C). The average value of transfection-negative cells in EB1/EB3 group is normalized to 100%. N = 15 cells. Shown as mean ± SD. Unpaired t tests were performed. (E) Percentage of HeLa cells with liquid-like condensates. Cells with high expression of Janusein or EB1 are analysed. The behavior of EB3 is similar to EB1. Chi-Square test was performed. ns, not significant; **** p < 0.0001.

**Figure S4.**
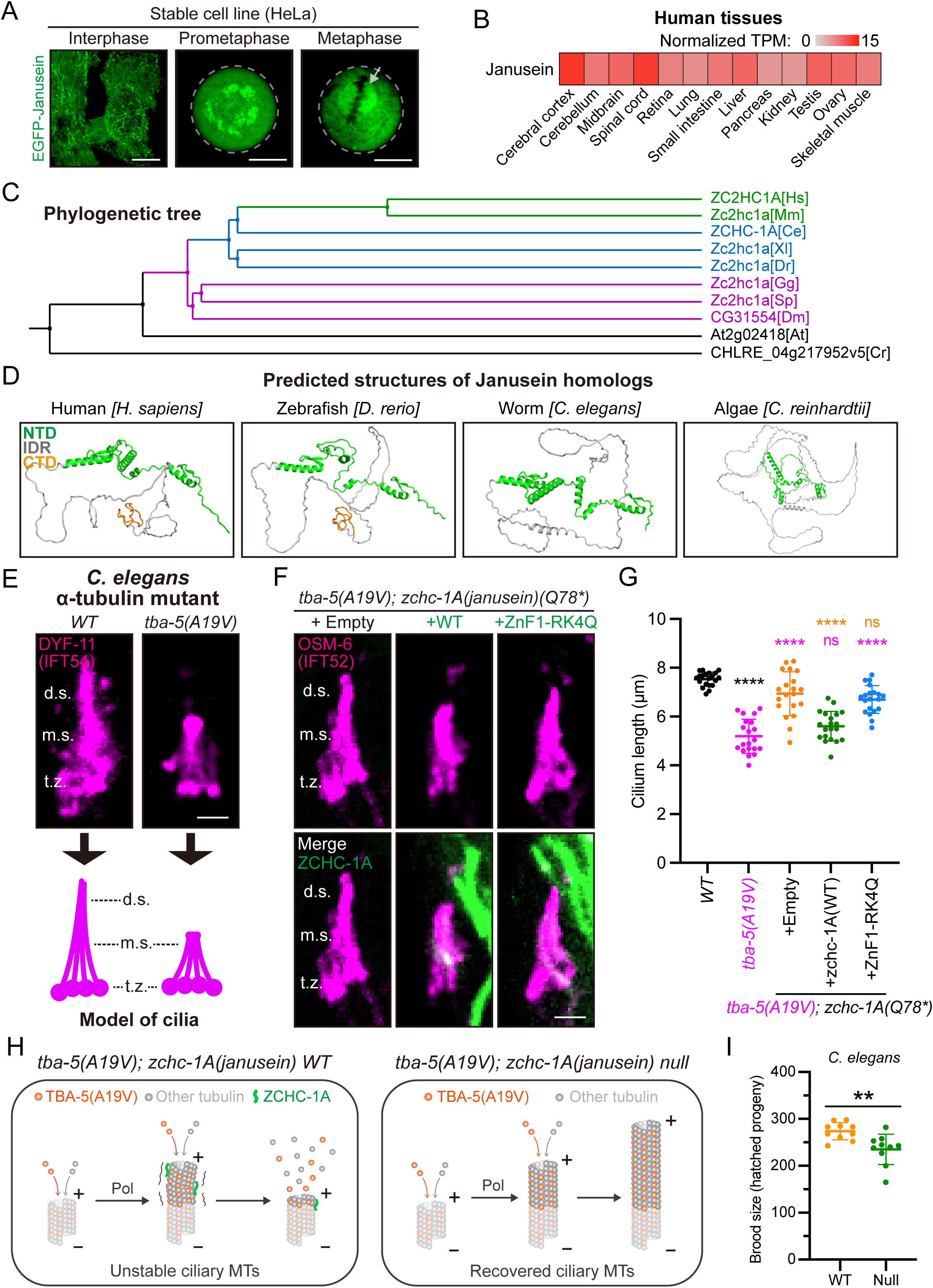
Janusein is conserved across eukaryotic species. (A) Representative live-cell images showing subcellular localizations of EGFP-Janusein in a stable HeLa cell line. (left) Interphase, (middle) prometaphase, (right) metaphase. The mitotic spindle is indicated by a white arrow. Scale bar, 10 μm. (B) Relative expression levels of endogenous Janusein (ZC2HC1A) across human tissues. Normalized TPM (transcripts per million mapped reads) values were obtained from Human Protein Atlas (https://www.proteinatlas.org). (C) Phylogenetic tree of Janusein (ZC2HC1A) homologs in representative eukaryotic species (see Methods). (D) AlphaFold3-predicted structures of Janusein homologs from four representative eukaryotic species in (C). (E) Representative images showing the morphology of amphid cilia in wild-type *Caenorhabditis elegans* and the α-tubulin mutant *tba-5 (A19V)*. DYF-11(IFT54)::mScarlet marks the cilia. Models of ciliary structures are shown below. t.z., transition zone; m.s., middle segment; d.s., distal segment. Scale bar, 2 μm. (F) Representative cilia images in *tba-5 (A19V)*; *zchc-1A (Janusein homolog) (Q78*)* animals overexpressing empty vectors (left), or wild-type P*zchc-1A::zchc-1A::gfp* (middle), or the P*zchc-1A::zchc-1A::gfp (ZnF1-RK4Q)* mutant. OSM-6(IFT52)::mScarlet marks the cilia. Scale bar, 2 μm. (G) Quantification of ciliary length in each group in (E and F). N = 20 amphids. Shown as mean ± SD. One-way ANOVA with Tukey’s tests were performed. ns, not significant; **** p < 0.0001. (H) (Left panel) Schematic model of α-tubulin TBA-5 (A19V) disrupting ciliary MTs in *C. elegans*. The ‘gain-of-function’ TBA-5 (A19V) continuously assembles into ciliary MTs and destabilizes them in the presence of wild-type endogenous ZCHC-1A. (Right panel) Schematic model of ZCHC-1A null allele rescues the ciliary defects induced by TBA-5 (A19V). Loss of ZCHC-1A is presumed to result in increased MT growth, thus recovering ciliary MTs. Pol, polymerization. (I) Quantification of *C. elegans* brood size in wild-type (N2) or *zchc-1A (Q78*)* (Null) animals. N = 10 animals. Shown as mean ± SD. Unpaired t tests were performed. ** p < 0.01.

**Figure S5.**
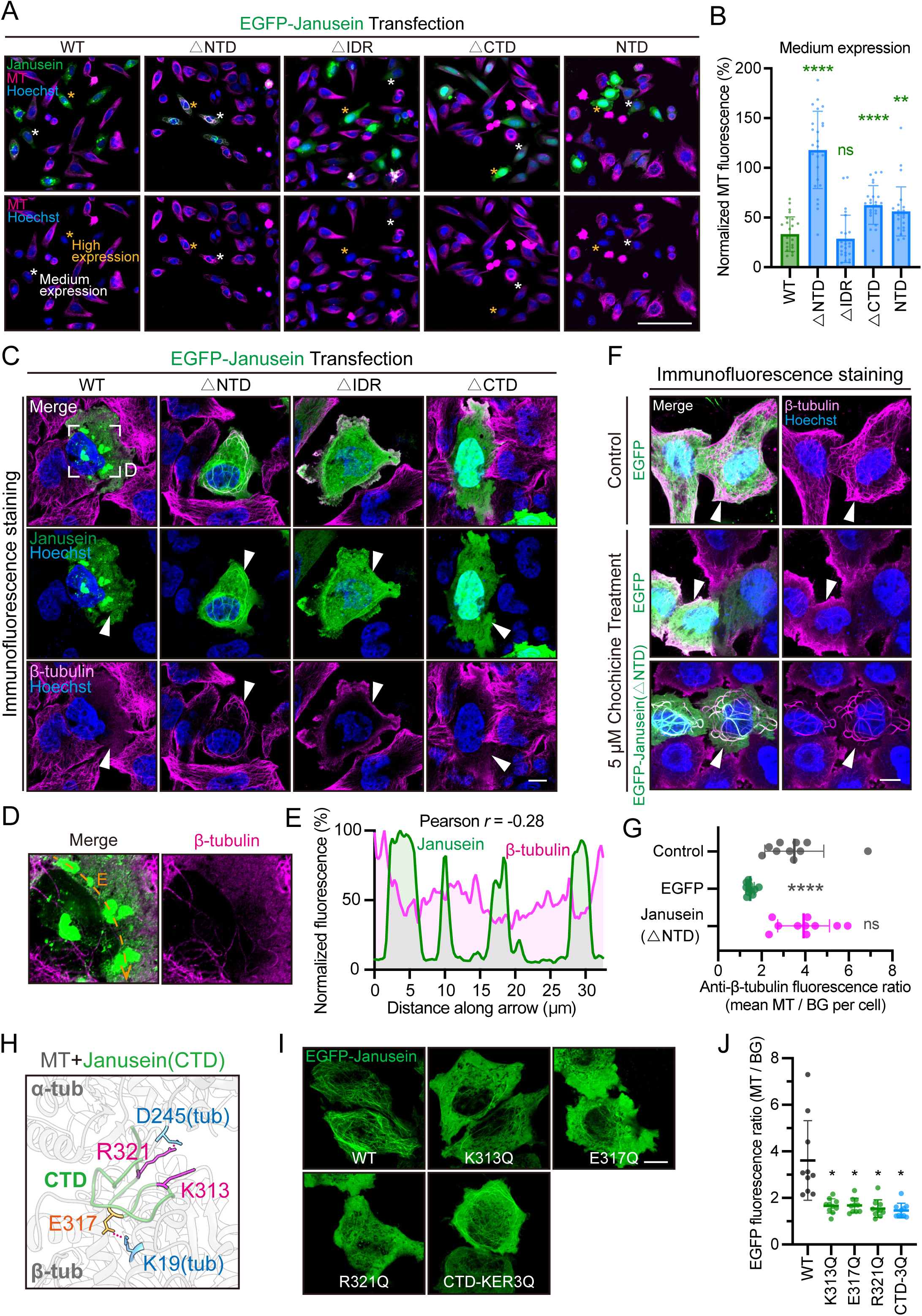
The C-terminal fragment of Janusein binds and stabilizes MTs. (A) Low-magnification overview of Janusein truncation mutants-transfected HeLa cells. Representative cells with medium (white) or high (orange) transfection are indicated by asterisks. Scale bar, 100 μm. (B) Whole-cell MT fluorescence intensities in HeLa cells with medium expression of Janusein truncation mutants. For each group, the average MT fluorescence of 10 randomly selected transfection-negative cells is normalized to 100%. Shown as mean ± SD. N = 25 cells. Welch’s ANOVA with Dunnett’s tests were performed. ns, not significant; ** p < 0.01; **** p < 0.0001. (C) Representative immunofluorescence images of HeLa cells with high expression of EGFP-Janusein truncation mutants. Transfection-positive cells are indicated by white arrowheads. Scale bar, 10 μm. (D) Enlarged view of the wild-type Janusein-expressing cell in (C). (E) Fluorescence intensity profiles of Janusein (green) and β-tubulin (magenta) along the arrow. The maximal fluorescence for each group is normalized to 100%. (F) Representative immunofluorescence images of HeLa cells with or without colchicine (5 μM) treatment. Transfection-positive cells are indicated by arrowheads. Scale bar, 10 μm. (G) Quantification of anti-β-tubulin fluorescence ratios of MTs to their adjacent backgrounds (BG) for each group in (F). For each cell, the average ratio of 10 independent MTs is calculated. N = 10 cells. Shown as mean ± SD. Welch’s ANOVA with Dunnett’s tests were performed. (H) Predicted interaction pattern of the Janusein C-terminal domain (CTD) with the MT inner surface of human tubulins (TUBA4A/TUBB8). Key interacting residues are highlighted in magenta and orange (Janusein), or blue (tubulins). Salt bridges are indicated by dotted lines. (I) Representative live-cell images of HeLa cells with high expression of Janusein CTD mutants. Scale bar, 10 μm. (J) Quantification of EGFP-Janusein fluorescence ratios of MTs to their adjacent backgrounds (BG) for each group. For each cell, the average ratio of 10 independent MTs is calculated. N = 10 cells. Shown as mean ± SD. Welch’s ANOVA with Dunnett’s tests were performed. ns, not significant; * p < 0.05; ** p < 0.01; *** p < 0.001; **** p < 0.0001.

**Figure S6.**
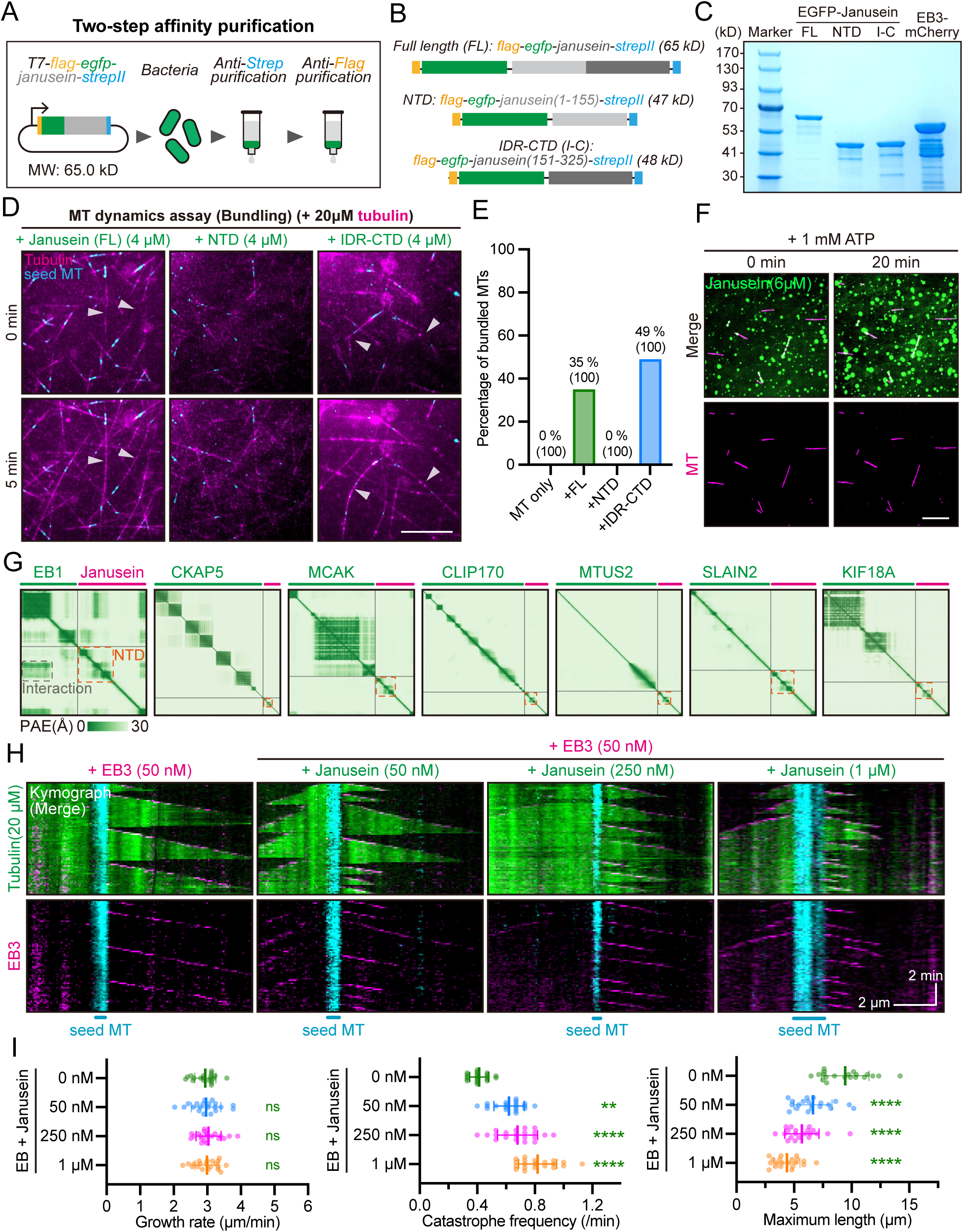
Janusein alone promotes MT bundling in vitro. (A) Schematic model showing two-step affinity purification of Flag-EGFP-Janusein-StrepII from bacteria. MW, molecular weight. (B) Schematic showing EGFP-Janusein and its truncation mutants for purification. (C) SDS-PAGE analysis of purified EGFP-Janusein and EB3-mCherry from bacteria. (D) Representative TIRF images showing in vitro MT-bundling activity of Janusein. White arrowheads indicate sites where two MTs bundle together. Scale bar, 10 μm. (E) Percentage of bundled MTs in each group in (D) after 10-min MT polymerization. N = 100 MTs per group. (F) Representative images showing in vitro co-incubation of MTs and Janusein, in the presence of 1 mM ATP. Scale bar, 5 μm. (G) AlphaFold3-predicted interaction patterns between Janusein (magenta) and MT plus-end-binding proteins (green), including EB1, CKAP5, MCAK, CLIP170, MTUS2, SLAIN2 and KIF18A. Shown as predicted alignment error (PAE) plots with expected position error (Å) ranges from 0 (green) to 30 (white). The NTD of Janusein is indicated by orange boxes. (H) Representative kymographs displaying the time-lapse dynamics of growing MTs. HiLyte647-conjugated seed MTs (cyan), HiLyte488-conjugated tubulins (green), EGFP-Janusein (green), and EB3-mCherry (magenta) were employed. Scale bar, 2 μm (horizontal) and 2 min (vertical). (I) MT growth rate (left), catastrophe frequency (middle), and maximal growth length (right) for each group in (H). One-way ANOVA with Dunnett’s tests (left, right), or Kruskal-Wallis test with Dunn’s multiple comparisons (middle) were performed. ns, not significant; ** p < 0.01; **** p < 0.0001.

**Figure S7.**
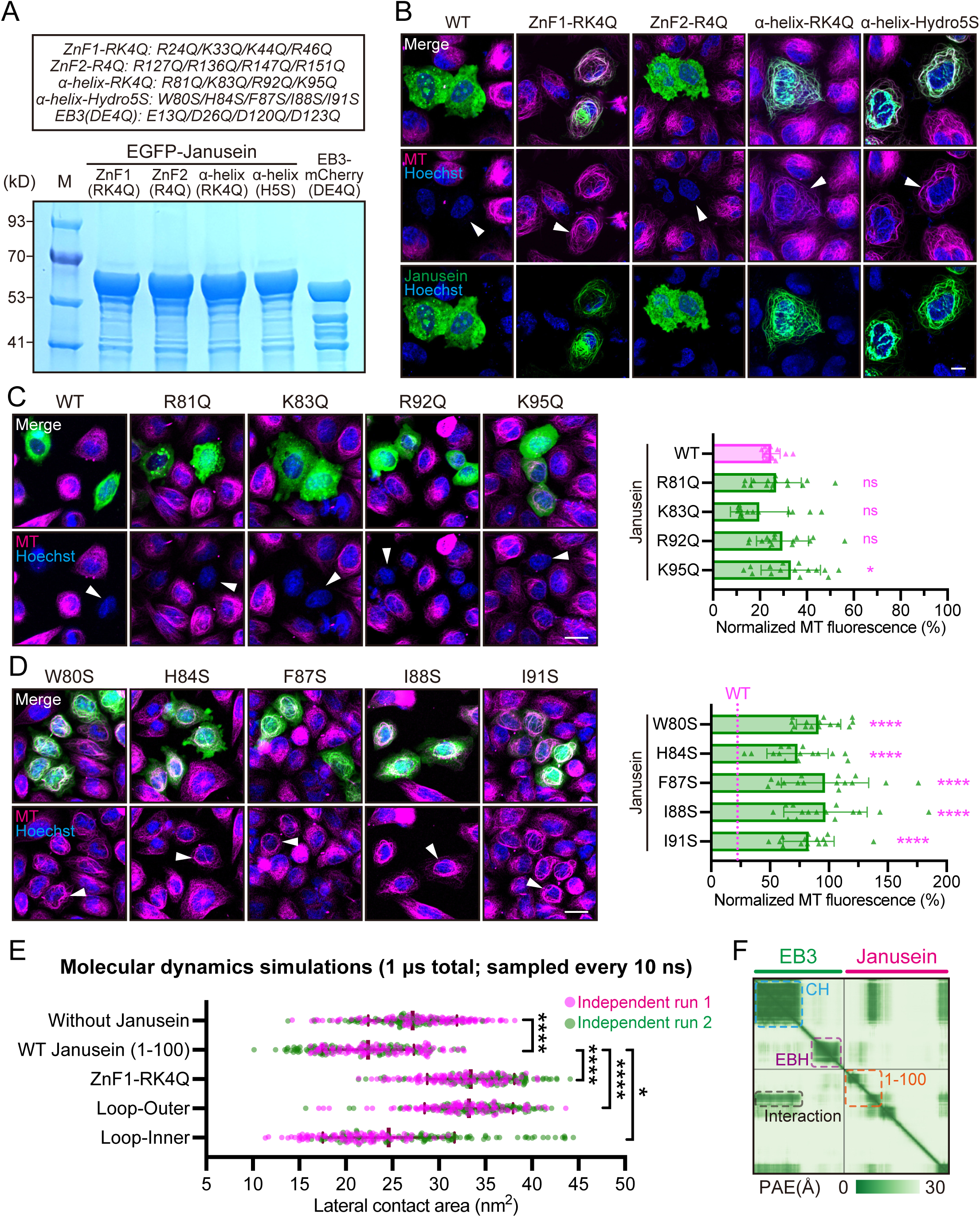
The Janusein NTD interacts with MT protofilament for destabilization. (A) SDS-PAGE analysis of EGFP-Janusein mutants and the EB3 (DE4Q) mutant purified from bacteria. M, marker. (B) Representative live-cell images of HeLa cells with high expression of Janusein mutants, relates to Figure 5I. Transfection-positive cells are indicated by white arrowheads. Tubulin Tracker Deep Red (magenta) was used to stain MT networks. Scale bar, 10 μm. (C) (Left) Representative live-cell images of HeLa cells with high expression of Janusein single mutants within the α-helix (electropositive residues). Transfection-positive cells are indicated by white arrowheads. Scale bar, 20 μm. (Right) Statistics of normalized MT fluorescence in transfection-positive cells. Control group in Figure 5I (transfection-negative group) is normalized to 100%. N = 15 cells. Shown as mean ± SD. Unpaired t tests were performed. (D) (Left) Representative live-cell images of HeLa cells with high expression of Janusein single mutants within the α-helix (hydrophobic residues). Transfection-positive cells are indicated by white arrowheads. Scale bar, 20 μm. (Right) Statistics of normalized MT fluorescence in transfection-positive cells. Control group in Figure 5I (transfection-negative group) is normalized to 100%. The mean value of WT group in (C) is highlighted by magenta dashed lines in (D). N = 15 cells. Shown as mean ± SD. Unpaired t tests were performed. (E) Statistics of lateral contact area of adjacent MT protofilaments in the absence or presence of Janusein mutants during molecular dynamics simulations. For each system, the total simulation time is 1,000 ns and sampling is conducted every 10 ns. Two independent simulations were performed for each group. Shown as mean ± SD. Kruskal-Wallis test with Dunn’s multiple comparisons were performed. ns, not significant; * p < 0.05; **** p < 0.0001. (F) AlphaFold3-predicted alignment error (PAE) plot of full-length EB3 (green) and Janusein (magenta). Expected position error (Å) ranges from 0 (green) to 30 (white). The EB3 calponin homology (CH) domain is indicated in blue, EB homology (EBH) domain in purple, Janusein (1-100) fragment in orange.

**Movie S1. Time-lapse live-cell imaging of HeLa cells over-expressing EGFP-EB1 and EB3-mCherry (left), or EGFP-Janusein and EB3-mCherry (right).** This movie corresponds to Figure 2A. Total time: 60 sec (12× sped up).

**Movie S2. Time-lapse TIRF imaging of in vitro microtubule dynamics with EB3 and Janusein.** This includes GMPCPP-stabilized seed MTs (cyan), porcine brain tubulins (green) (20 μM), EB3-mCherry (magenta) (50 nM), and EGFP-Janusein (green) (250 nM). This movie corresponds to Figure 4E. Total time: 10 min (120× sped up).

**Movie S3. Molecular dynamics simulations showing dynamics of the Janusein (1-100) fragment.** ZnF1 (blue), semi-flexible loop (orange) and α-helix (magenta) are differentially colored. This movie corresponds to Figure 6H. Total simulation time: 300 ns.

## Notes

### Competing Interest Statement

The authors have declared no competing interest.

